# A rigid anchor and a flexible blade: calcium activates gelsolin through an asymmetric conformational ensemble

**DOI:** 10.64898/2026.10.16.759396

**Authors:** Bruno Stegani, Loic Girois, Federico Ballabio, Toni Giorgino, Riccardo Capelli, Guillaume Gerbaud, Michela Bollati, Mario Milani, Alessio Bonucci, Carlo Camilloni, Matteo de Rosa

## Abstract

Gelsolin is the founding member of a superfamily of multidomain proteins that remodel the actin cytoskeleton in response to calcium. How the full-length protein is activated remains unclear because its active form has escaped structural characterization. Here, we combine small-angle X-ray scattering, pulsed EPR spectroscopy and molecular simulations to follow its transition from the compact inactive state to the active form. Activation proceeds through sequential, domain-specific rearrangements and culminates not in a single structure but in a heterogeneous ensemble, in which a comparatively rigid half implicated in filament capping coexists with a highly dynamic half implicated in severing. These states are populated progressively across the nanomolar-to-millimolar calcium concentrations that gelsolin encounters inside and outside cells, linking local calcium availability to the protein’s functional repertoire. Gelsolin activation thus emerges as a graded population shift rather than a switch between discrete conformations, providing a structural framework for its diverse functions and, more generally, for the regulation of dynamic multidomain proteins.

## Introduction

Actin is one of the most abundant proteins in eukaryotic cells. It participates in many physiological processes, and its dynamics are highly regulated by members of the gelsolin-like protein superfamily (InterPRO IPR036180), which includes eight proteins in humans: gelsolin, adseverin, villin-1, advillin, supervillin, villin-like protein, flightless-1 homolog and CapG. Gelsolin (GSN) is the founding member of the superfamily; it was first isolated in 1979 and named after its ability to modulate the gel-sol transition of cytoplasmic actin (1). GSN is now known to possess a variety of actin regulatory functions: it can sever, cap, and nucleate actin filaments, and sequester monomers (reviewed in (2)). These actin regulating activities impact biological functions as varied as the processes in which actin participates, including cell motility, ion channel regulation, cell signaling, apoptosis, phagocytosis, secretion, platelet activation and modulation of phospholipase C activity (reviewed in (3)).

Over time, additional putative functions have been assigned to GSN-like proteins, including roles in nuclear hormone receptor-mediated signaling, wound repair, cell morphology, invasion and migration, epithelial-to-mesenchymal transition, and regulation of podosomes and cadherin junctions (4–6). In parallel with the centrality of these biological processes, GSN has been associated with a wide range of diseases: for example, increased expression has been detected in idiopathic interstitial pneumonia, cardiac diseases and oxidative stress, while its downregulation has been observed in various types of cancer and in rheumatoid arthritis (reviewed in (7)). Yet, the exact molecular role of GSN in these pathological conditions has not been fully clarified. An exception is the identification of a physical interaction between GSN and human papillomavirus oncoproteins (8, 9), and the recent discovery of a key role for GSN in neoplastic cell dormancy (10), making the protein a novel promising antitumor target. Several pathogenic GSN substitutions associated with the rare monogenic disease AGel amyloidosis have also been extensively characterized, showing mutation-specific molecular mechanisms and clinical outcomes (11–13).

Three different isoforms of GSN have been identified in humans, produced through alternative splicing or transcription initiation from the same gene. Isoform 3 is preferentially expressed in the central nervous system (14). The other two forms, which are almost ubiquitous, are referred to as the cytoplasmic and plasma (secreted) proteins, respectively. The mature (signal peptide-cleaved) form of the plasma isoform is hereafter used as the reference sequence. The two ubiquitous isoforms experience very different calcium environments. Cytoplasmic gelsolin is exposed to a resting free Ca²⁺ concentration of ∼100 nM, to global transients that rarely exceed ∼1 µM, and to local microdomains near open channels where free Ca²⁺ can transiently reach tens of micromolar (15). Plasma gelsolin is synthesized along the secretory pathway, where luminal free Ca²⁺ ranges from ∼100 µM in the trans-Golgi to several hundred micromolar in the endoplasmic reticulum (16), and then circulates at ∼2–3 µM in plasma, where ionized Ca²⁺ is ∼1.2 mM. In plasma, gelsolin and Gc-globulin form the extracellular actin-scavenger system that clears actin released from damaged tissue (17); gelsolin depletion in sepsis, trauma and burns predicts adverse outcomes, which has motivated the clinical development of recombinant plasma gelsolin (18).

All GSN-like members are modular and multifunctional proteins. GSN contains six prototypical gelsolin-like domain (named G1 to G6), each composed of 97-118 residues. Six such domains are also present in the homologues, except for CapG, which contains only three domains. Each G domain contains a calcium-binding site (type-2); in addition, GSN harbors two incomplete Ca²⁺ sites (type-1), located at the actin interfaces in G1 and G4 (19). In the absence of calcium, GSN adopts a closed conformation with the G1 to G5 domains wrapped around G6, burying all actin-binding surfaces; this state is also referred to as the inactive form. Binding of calcium ions promotes conformational changes that lead to an open structure with the actin-binding sites solvent-exposed (20). Calcium is not the only modulator of GSN conformation and activity: several other molecules and environmental factors act synergistically or competitively. A key role in the GSN life-cycle is played by phosphatidylinositol phosphates (PIPs), both bis- and tris-phosphorylated. GSN is predicted to contain two PIP-binding sites, and their binding promotes protein inactivation even at high Ca²⁺ concentrations (21–23). ATP also competes for at least one of the PIP-binding sites and exerts a similar regulatory effect (24, 25). Beyond ligand control, temperature and pH can also activate GSN in a Ca²⁺-independent manner (22, 26). Although these conditions fall outside physiological ranges, they likely contribute to GSN behavior in complex intra- and extracellular environments. In addition, gelsolin can be irreversibly activated by caspase-3 cleavage of the long linker between G3 and G4 (27). This mechanism is relevant during apoptosis, when the cytoskeleton must be dismantled and the cell cleared.

Early studies (28, 29) showed that Ca²⁺ -induces large but reversible conformational changes, occurring through sequential events and involving one or more intermediate states. However, these studies disagreed on both the number of events and the Ca²⁺ concentrations required to trigger them. Functional titrations are similarly stratified: full-length gelsolin binds F-actin with half-maximal effect at ∼0.1 µM free Ca²⁺ and exposes limited proteolysis sites and a first actin-binding site between 0.3 and 1 µM, whereas efficient severing and nucleation at neutral pH require ∼10 µM Ca²⁺; tryptophan fluorescence resolves three transitions with apparent Kd values of ∼0.1, 0.3 and 6 µM, the last associated with release of the C-terminal tail latch (30–33). How this stratified calcium dependence relates to the global conformational transition has not been established. Later structural investigations, including high-resolution crystallographic structures of equine and human full-length GSN (19, 34), enabled the identification of all eight Ca²⁺-binding sites and set the basis for subsequent mechanistic studies. Additional biophysical analyses (20, 35) described activation as a two-step, three-state process that not only shifts the protein toward a more extended shape but also dramatically increases its conformational flexibility, likely explaining why high-resolution structures of the active state have never been obtained. Crystal structures of Ca²⁺-bound fragments of GSN are available (34, 36–39), revealing subtle local Ca²⁺-induced rearrangements and latches that must be released to prevent the protein from remaining locked in its closed conformation.

The recently reported (40) cryo-EM structure of the GSN:F-actin capping complex showed that GSN adopts an extended conformation in which the G4-G6 triplet remains compact, while G1 extends to bind the filament tip, sterically preventing repolymerization. Based on this structure, the authors propose a severing mechanism in which G1 plays a major role by bending and displacing the filament. The same group further investigated the molecular basis of severing and showed that the isolated G2-G3 fragment binds F-actin in a stepwise manner, inducing local conformational changes that may later facilitate recruitment of the other domains (41).

Despite these significant advances, neither the structure of fully active GSN nor that of the likely transient severing complex are available. Moreover, the sequence of conformational events linking Ca²⁺ binding to domain exposure and filament engagement remains incompletely understood, and it is unclear how the calcium dependence of activation relates to the very different calcium environments in which the cytoplasmic and plasma isoforms operate. The present study contributes to addressing these questions by characterizing the activation pathway of full-length gelsolin and the conformational ensemble populated by its active state, and by mapping both onto the functional and cellular calcium scales.

## Materials and Methods

Please refer to the *supplementary information* for detailed explanation of all Materials and Methods, here only briefly summarized.

### Protein production

Human full-length gelsolin and engineered double-cysteine variants were recombinantly expressed in *E. coli* and purified by affinity, ion-exchange, and size-exclusion chromatography as reported in (11).

### SAXS measurement

Small-angle X-ray scattering (SAXS) data were collected on purified gelsolin at multiple protein and Ca²⁺ concentrations. Scattering profiles were analyzed using standard procedures to determine molecular weight, radius of gyration (R_g_), and pair-distance distribution functions.

### 4p-DEER (Q-band) Spectroscopy

To monitor Ca²⁺-induced interdomain rearrangements, seven double-cysteine gelsolin variants were generated by site-directed mutagenesis, spin-labeled with MTSL, and analyzed by Q-band four-pulse DEER spectroscopy. Distance distributions were extracted from background-corrected traces using Tikhonov regularization.

### Molecular Dynamics and integrative modelling

The conformational landscape of Ca²⁺-activated gelsolin was investigated using a multi-eGO (42, 43) structure-based model parameterized from all-atom molecular dynamics simulations of the closed full-length protein and of calcium-bound domains. Independent simulations were performed to generate ensembles of activated conformations and to characterize opening pathways.

Agreement with experimental data was achieved through an integrative modeling strategy combining SAXS and DEER-EPR measurements with Bayesian Maximum Entropy (BME) reweighting (44). The resulting ensemble was further analyzed using solvent-accessible surface area calculations, Markov state models (MSM), and Energy Landscape Visualization Method (ELViM) (45–48) projections to characterize conformational states, domain rearrangements, and activation pathways. Additional opening simulations were used to determine the temporal hierarchy of domain exposure during activation.

### Use of generative AI

ChatGPT 5.6 was used to improve the readability and linguistic quality of the manuscript. All generated suggestions were reviewed and edited by the authors, who take full responsibility for the final content of the manuscript.

## Data, Materials, and Software Availability

SAXS datasets have been deposited in SASBDB under accession numbers SASD2Z8–SASD2R9 (see Table S1). Computational data generated in this study are available in Zenodo. Simulation data, analysis scripts, and software used in this study are available in DOI: 10.5281/zenodo.23237877. Opening simulations, activation-pathway analyses, and associated scripts are available in DOI: 10.5281/zenodo.23235294. The final reweighted ensemble of activated gelsolin, including simulation trajectories, statistical weights, and representative PDB structures for each conformational state, has been deposited in DOI: 10.5281/zenodo.23237909.

## Results

### Cooperative Ca²⁺-binding drives asymmetric opening of gelsolin

Small angle X-ray scattering (SAXS) is a well-established technique to investigate the conformational landscape of GSN. Here it was used for an initial evaluation of the dependence of protein conformation on Ca²⁺ concentration through batch measurements collected at different protein (from 2.7 to 52.1 μM) and Ca²⁺ (0-10 mM) concentrations. Under all tested conditions, the molecular weight estimated either by Bayesian analysis (Mw(B) = 80.6 kDa, PRIMUS) or from the absolute scattering intensity (Mw(abs) = 75.9 kDa) was consistent with a monomeric species, excluding oligomerization events throughout the investigated concentration range.

The closed conformation (1 mM EDTA) displayed a constant radius of gyration (Rg) across all protein concentrations tested (R_g_ = 31.7 ± 0.2 Å), whereas the fully activated state (10 mM Ca²⁺) exhibited a markedly larger radius of gyration and slightly greater variability (R_g_ = 40.8 ± 0.8 Å). At all protein concentrations investigated, GSN appeared to have completed the global opening transition at Ca²⁺ concentrations above ∼200 μM (Table S1 and Figure S1).

The SAXS measurements provide an effective description of the calcium-dependent opening of GSN. Here, we used SAXS to investigate how the global conformation of GSN depends on both protein and Ca²⁺ concentrations. Plotting the R_g_ as a function of GSN concentration at a fixed total Ca²⁺ concentration of 50 μM revealed a smooth transition between the closed and open conformations (Figure 1A), Rg decreases as GSN concentration increases consistent with progressive depletion of free Ca²⁺ through binding to GSN, favouring the closed conformation. To estimate the half-transition free Ca²⁺ concentration (K_1/2_), we fitted the data using a simple two-state model with an effective ion-binding occupancy (n_eff_; see Methods 4). The fit yielded n_eff_ = 6.4 ± 0.9 and K_1/2_ = 18 ± 4 μM.

**Figure 1:**
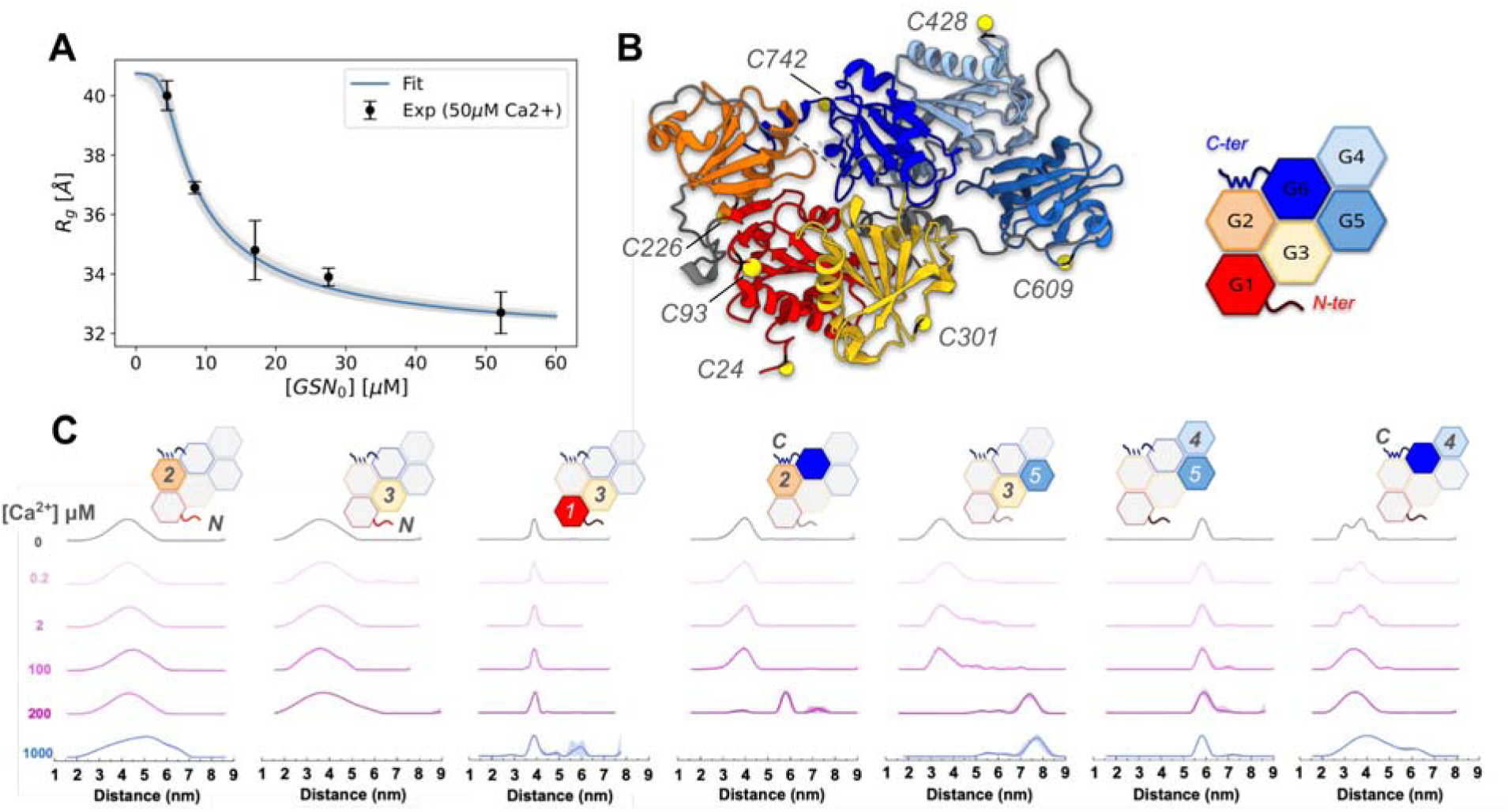
**A)** Variation of the gyration radius (R_g_) measured at a fixed Ca²⁺ concentration of 50 μM as a function of GSN concentration. Experimental data (black) were fitted using a two-states model that account for Ca²⁺ depletion (cyan curve) resulting in a half-transition free Ca²⁺ concentration K_1/2_ = 18 ± 4μM and an effective ion-binding occupancy n_eff_ = 6.4 ± 0.9. **B)** cartoon and schematic representation of GSN in the closed (Ca²⁺ -free) conformation. Domains are color-coded and labeled throughout the manuscript, labeled cysteines are mapped onto the atomic model C) Distance distributions derived by background corrected Q-band DEER in the absence and in the presence of increasing concentration of Ca^2+^ obtained for the spin labelled GSN mutants bearing the tags mapped in panel B of this figure (Ca-free GSN (PDB: 3FFN).

Because GSN binds multiple Ca²⁺ ions, the free Ca²⁺ concentration can differ substantially from the total Ca²⁺ concentration, particularly at higher protein concentrations. The same model with the parameter obtained above was able to describe the R_g_ measurement across the full range of GSN and total Ca²⁺ concentrations (Figure S1).

The fitted n_eff_ is consistent with the known calcium-binding architecture of GSN, which contains six type-II Ca²⁺-binding sites; the two type-I sites may become relevant primarily in the presence of actin (37, 39). However, n_eff_ should be interpreted as an effective parameter rather than a direct measure of calcium-binding stoichiometry. Because GSN opening involves multiple domain rearrangements, the two-state model provides a useful description of the global transition without over-interpreting the SAXS dataset in terms of a detailed microscopic mechanism.

To gain a more detailed view of the activation process, we employed 4p-DEER measurements on a panel of engineered double-cysteine variants reporting inter-domain distances throughout the protein (Figure 1B). In the absence of Ca²⁺, all labeled GSN variants produced distance distributions (Fig. 1C) in excellent agreement with those calculated from the crystal structure of calcium-free GSN (PDB: 3FFN) (Fig. S2), validating the use of this approach to investigate conformational states that remain inaccessible to high-resolution structural methods.

DEER analysis revealed a hierarchical nature of the conformational changes accompanying activation. The largest Ca²⁺-induced structural rearrangement was observed for the G3-G5 reporter. Increasing calcium concentration progressively altered the dipolar frequency of the DEER oscillations, shifting the distance distribution maximum from 3.5 ± 0.4 nm in the absence of calcium to 7.2 ± 0.5 nm at saturating conditions (1 mM Ca²⁺) (Fig. 1C). This result indicates a marked separation between the N- and C-terminal halves of the protein upon activation. Limitations imposed by the phase-memory time (Tm) and instrumental settings prevented acquisition of the full oscillation period at saturating calcium (Fig. S3). The presence of longer-distance conformations, such as those observed in the F-actin:GSN capping complex (roughly 10 nm), cannot be excluded.

Interestingly, the G3-G5 reporter also revealed the possible presence of intermediate conformational states. DEER traces collected between 0.2 and 2 µM Ca²⁺ exhibited oscillation frequencies distinct from those observed in both apo and saturating conditions (Fig. S3), resulting in bimodal distance distributions with minor populations centered between 4 and 6 nm (Fig. 1C). At 200 µM Ca²⁺, the DEER trace and corresponding distance profile became indistinguishable from those observed at saturating calcium concentrations, indicating that the inter-triplet opening transition is largely complete. A second reporter connecting the two major structural regions of the protein, G2-C_ter_, displayed a similar, although less pronounced, behavior. Increasing Ca²⁺ concentrations from 0 to 200 µM shifted the distance distribution maximum from 3.9 ± 0.4 nm to 5.6 ± 0.4 nm (Fig. 1C), indicating progressive displacement of the C-terminal region away from G2. Upon reaching 1 mM Ca²⁺, the DEER modulation depth decreased markedly (Fig. S3), consistent with either a shift towards distances beyond the detection limit or a highly heterogeneous ensemble with no preferential interspin distance.

Analysis of reporters located within the G1-G3 triplet further clarified the structural basis of these large-scale rearrangements. For the N_ter_-G3 construct, addition of 1 mM Ca²⁺ resulted in a dramatic reduction of the DEER modulation depth compared with the apo state (Fig. 1), consistent with either an increase of the inter-domain distance from 3.6 ± 0.6 nm to values above the experimentally accessible range or such a heterogeneous population that no preferential conformation can be detected. The latter interpretation is supported by measurement of the N_ter_-G2 reporter that displays modest Ca²⁺-dependent variations, but a concentration dependent broadening of the peak (Fig. 1C). A similar behavior was observed for the G1-G3 reporter, where saturating calcium strongly reduced the DEER trace intensity (Fig. S3), again suggesting separation of the two labels beyond 8 nm or increased conformational flexibility. However, residual oscillations remained detectable, yielding a distance distribution partially overlapping the one observed in the absence of calcium. Overall, DEER reporters within G1-G3 triplet consistently reveal increases in interdomain distances and the emergence of highly heterogeneous distance distributions at saturating calcium concentrations.

In contrast, DEER reporters within the G4-G6 triplet indicated limited structural rearrangements. The G4-G5 construct displayed essentially unchanged DEER traces and distance distributions in the absence and presence of saturating calcium (Fig. 1C and S3), suggesting that the relative organization of these domains is largely maintained throughout activation.

The only notable exception within this region involved the C-terminal tail. In the calcium-free state, the G4-C_ter_ reporter placed the C-terminal segment in proximity to G4, at approximately 3.7 ± 0.4 nm (Fig. 1C). Upon addition of 1 mM Ca²⁺, the corresponding DEER trace lost most of its oscillatory character (Fig. S3), a feature commonly associated with highly dynamic or disordered conformational ensembles. Accordingly, the distance distribution broadened, spanning approximately 1-7 nm. The observed broadening likely arises from the combination of structural rearrangements within the G4-G6 block and some increased conformational flexibility of the C-terminal tail, although to a lesser extent than that observed for the N-terminal triplet. This behavior is not apparent in the recently reported cryo-EM structure of the capping complex, whose calculated distance profile is considerably narrower than the experimentally observed DEER distribution (Figure S2).

Taken together, the DEER data indicate that Ca²⁺ activation is dominated by a large-scale separation of the G1-G3 and G4-G6 triplets, which corresponds to the conformational transition also observed by SAXS. In contrast, the triplets undergo asymmetric internal rearrangements: G4-G6 largely preserves its compact architecture, while the G1-G3 block acquires substantial conformational flexibility upon activation.

### The activation pathway is characterized by sequential domain exposure

To characterize the hierarchy of conformational changes during GSN activation, we performed 50 opening simulations using the multi-eGO model (See Methods 13) starting from the closed conformation with all type-II calcium-binding sites occupied.

For each simulation, we monitored the distances between the centroids of adjacent domain surfaces (see Methods 13). For each adjacent domain pair, we then constructed cumulative distribution functions (CDFs) of separation times, defined as the first time point at which the inter-domain distance reached a threshold value of 3 nm. Multi-eGO (and coarse-grained models in general) exhibit accelerated, non-realistic timescales due to the smoothing of the energy landscape, and should therefore be interpreted only relatively, not in absolute terms.

The cumulative distributions of separation times (Figure 2) reveal a well-defined sequence of opening events, schematically represented in Figure 2B using the hexagon code. The earliest event is the initial separation of the two major domain blocks (G1-G3 and G4-G6), with the G1-G6, G2-G6, and G3-G6 separations occurring on very similar timescales (7.5, 7.4, and 6.8 ns, respectively). This agrees with the Ca²⁺-dependent changes observed for G3-G5 in the DEER experiments (Figure 1C). Following block separation, domain G3 detaches from domain G1 (t = 35.6 ns). Finally, we observe the separation of G1– G2, followed on a similar timescale by the separation of G4–G6 (t = 48.6 ns and t = 53.2 ns, respectively).

**Figure 2:**
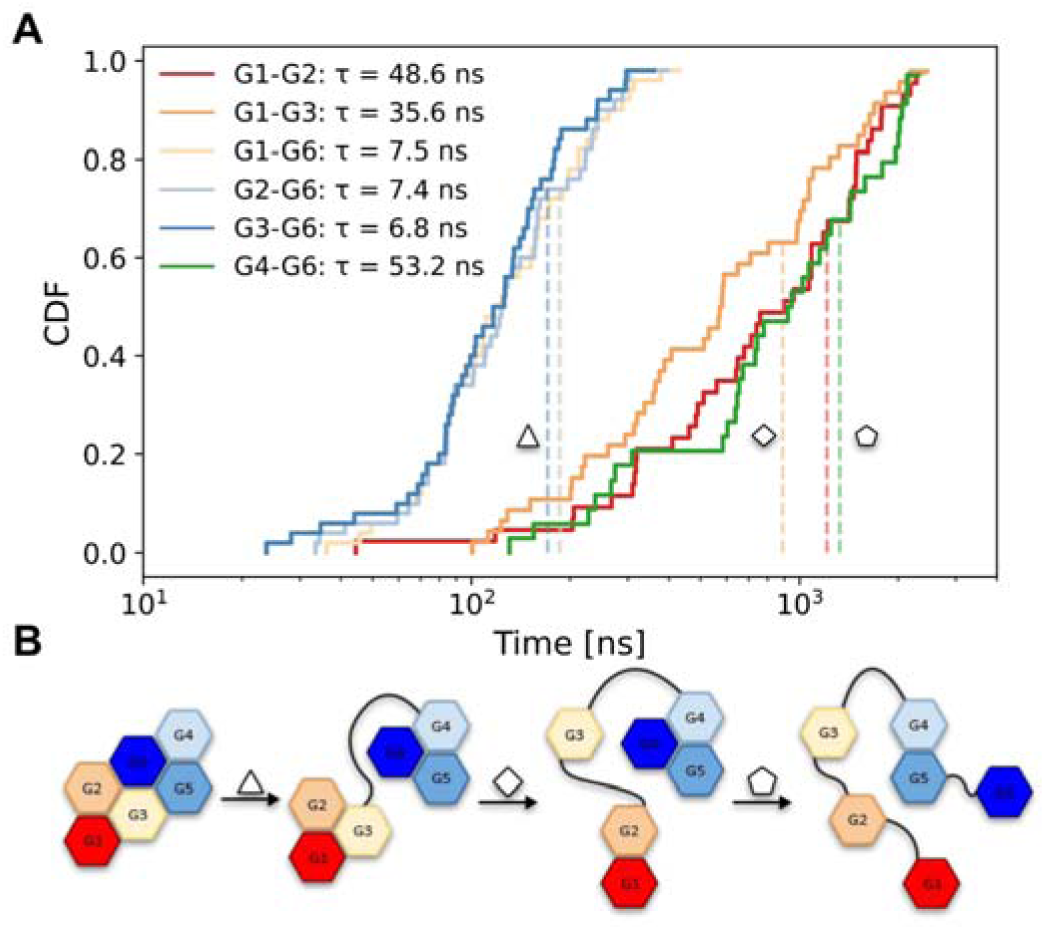
**A)** *In silico opening-time cumulative distribution functions (CDF) for each domain pair obtained from 50 independent opening simulation with type-II Ca^2+^ sites occupied.* Opening time was defined as the first time at which the distance between the interfaces of each domain-pair reaches 3 nm. **B)** *The resulting CDFs* reveal a domain-exposure hierarchy during gelsolin opening, in good agreement with the DEER measurements: first, separation of the G1–G2–G3 and G4–G5–G6 blocks; followed by detachment of G3 from G1–G2; and, finally, separation of G6 from G4–G5 and of G1 from G2. Note that the timescales of the simplified multi-eGO model are accelerated relative to real timescales and should therefore be interpreted comparatively rather than absolutely.

### Active gelsolin populates a heterogeneous conformational ensemble

To characterize the structural ensemble of calcium-activated gelsolin, we combined the multi-eGO simulation framework with SAXS and DEER measurements.

The multi-eGO model was parameterized using structural information extracted from all-atom molecular dynamics simulations (Methods 5) and designed to satisfy three main requirements: (i) preservation of the native fold of each individual domain, (ii) stable coordination of calcium ions at saturating concentrations, (iii) stabilization of an open-state ensemble capable of sampling the experimentally observed domain rearrangements associated with gelsolin activation (Methods 6). In addition, information for the full-length closed state was derived from previously reported simulations (49).

The resulting conformational ensemble was generated from three independent 15 µs simulations and subsequently reweighted using the Bayesian Maximum Entropy (BME)(44) framework to maximize consistency with experimental observations (Methods 8). Reweighting against the SAXS profile shifted the ensemble toward more extended conformations (Figure S9), yielding good agreement with most of the experimentally measured DEER signals. DEER distance distributions were back calculated using DEERPREdict (50) (Methods 9).

A notable exception was the G3-G5 distance (residues 301-609), for which the experimental DEER trace exhibited a substantial modulation depth (λ ⍰ 0.15) but no evidence of the short-distance population, present instead in the simulated ensemble. To account for this discrepancy, we exploited the Bayesian formulation of the reweighting procedure by introducing a prior based on the G3-G5 DEER data (Methods 9). Starting from this DEER-informed prior, we subsequently optimized agreement with the SAXS profile using BME, obtaining excellent agreement with the experimental scattering data (χ2 ≈ 1 for all replicas) using a regularization parameter θ = 100 (Figure S8).

The final ensemble, defined by the combination of simulation frames and their corresponding BME weights, reveals that calcium-bound gelsolin occupies a broad conformational landscape intermediate between the closed and actin-capping states. This behavior is evident from the distribution of R_g_ (Figure 3A), which spans values ranging from those of the closed structure (PDB: 3FFN) to those of the actin-capping conformation (PDB: 8VIZ, actin removed).

**Figure 3:**
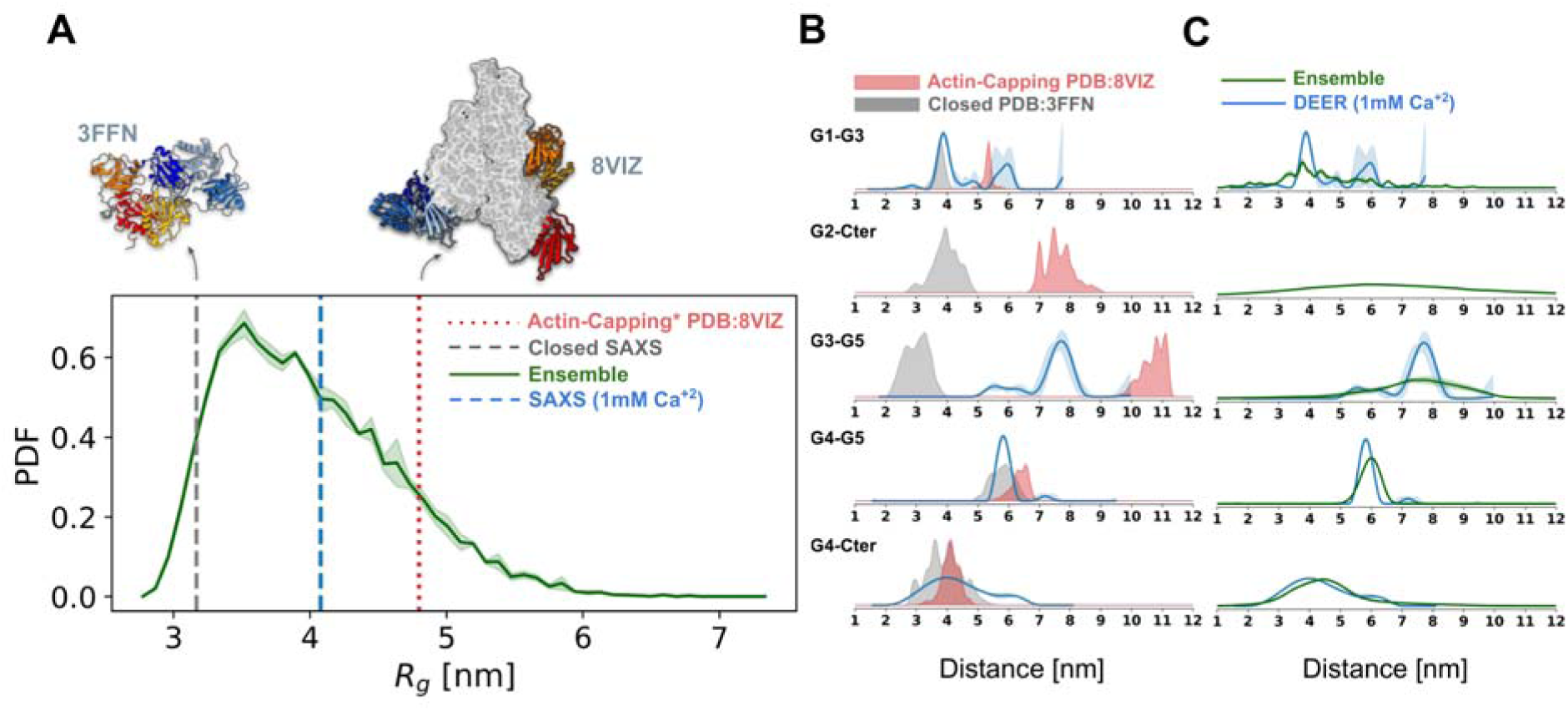
**A)** Radius of gyration (R_g_) distribution of the final reweighting ensemble. As a reference, R_g_ values are shown for GSN in the absence of Ca^+2^ (experimental average SAXS R in gray dashed lines) and for the structure of the actin-capping complex (PDB:8VIZ, red dotted line). The computed Rg value from the PDB of the Actin-capping structure does not account for the N-terminal HIS-tag present in all other measurements, as indicated by the “*” superscript in legend. Reference average experimental R_g_ of Gelsolin in the open state (at saturating concentration of ions, i.e. 1mM) is marked by a blue dashed vertical line. **B)** DEER distances (blue) of fully activated GSN (1 mM calcium) compared to spin-spin distribution computed with MMM2019 (51) obtained respectively for the closed (grey) and the actin capping (red) states of GSN. In-silico distance distributions have been computed from the available model of the full-length system for its closed (PDB: 3FFN) and actin-capping (PDB: 8VIZ) conformations. **C)** Comparison between experimental DEER distances (Blue) of fully activated GSN (1 mM Ca^2+^) and DEER distance distributions obtained from the open state Ensemble (Green). G2-C_ter_ experimental distance distribution is not shown due to the lack of clear oscillation in the time signal and hence the difficulty in deconvolving the signal into a distance distribution.

The resulting ensemble is consistent not only with the experimental R_g_ distribution, but also with the experimental DEER distance distribution (Figure 3C). The G3-G5 DEER distance, identified as the main reporter of the opening, populates under calcium-saturating conditions values intermediate between those characteristic of the closed and actin-capping states (Figure 3B). All other reporters further indicate that the activated state of GSN differs from the actin-capping complex both in relative distances and in the extent of its conformational flexibility (Figure 3B, C).

Analysis of SASA further supported this picture (Methods 10). For most domains, the exposure of actin-interacting surfaces lies between those observed in the closed and actin-bound structures, indicating partial activation. The main exception is domain G2 (Fig. S14:G2), whose actin-binding surface is predominantly exposed, suggesting that this domain adopts an actin-capping-competent configuration in a large fraction of the ensemble.

To visualize and characterize the conformational heterogeneity of the open state, we projected the reweighted ensemble onto the Energy Landscape Visualization Method (ELViM) (45, 46) coordinates (Methods 11). In this representation, nearby points correspond to structurally similar conformations, while population density reflects equilibrium free-energy basins. The resulting landscape (Figure 4) reveals a highly heterogeneous ensemble with multiple populated states. The most populated regions are located along the periphery of the projection and correspond to conformations with large R_g_ (Figure 4). Inspection of representative structures from these basins indicates that domains G1, G2, and G3 adopt highly variable relative orientations, whereas the G4-G5-G6 block remains comparatively compact and resembles its closed-state organization.

**Figure 4.**
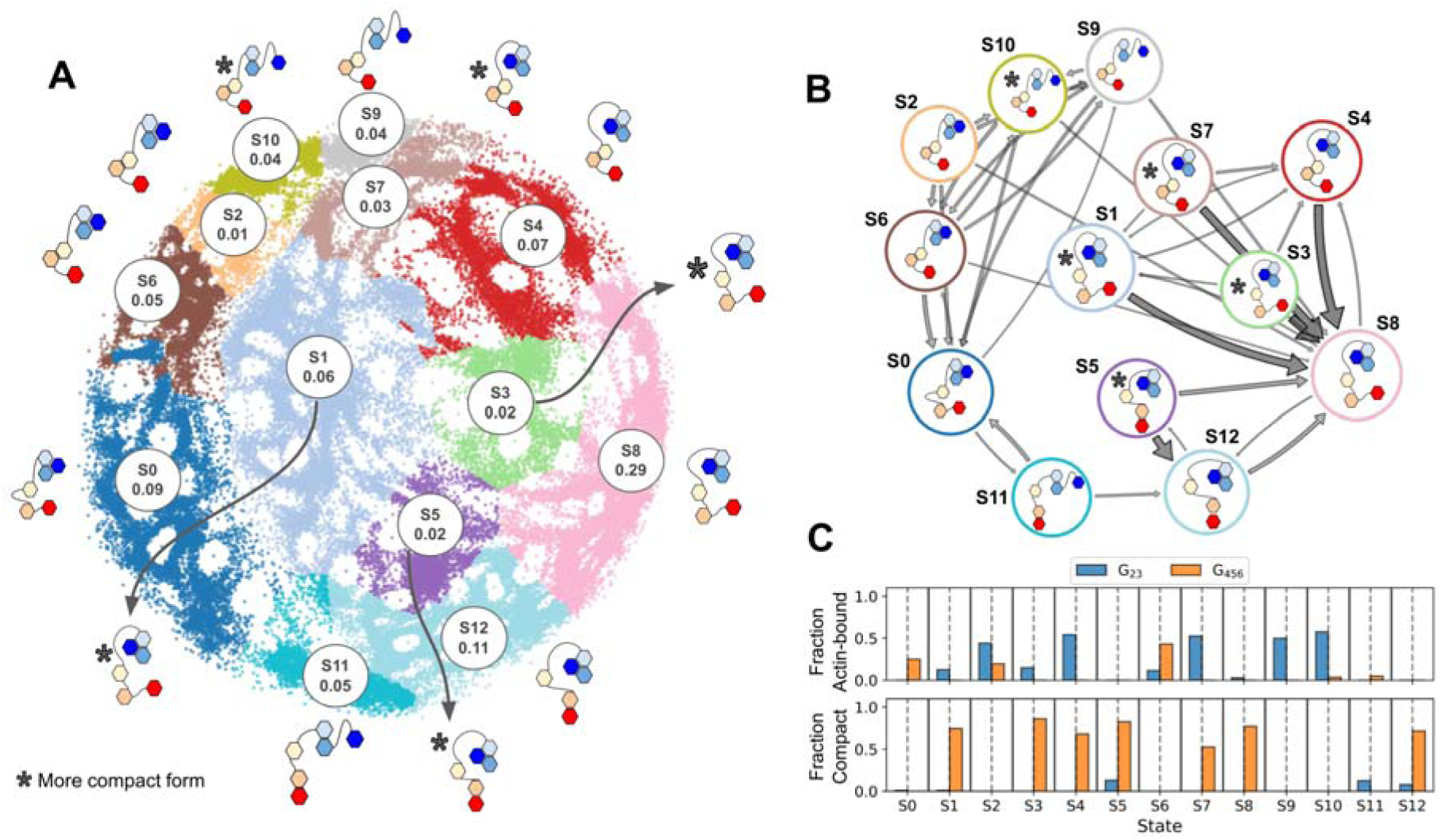
The ensemble is projected onto a 2-dimensional surface using an ELViM projection. The projection is used to define interpretable states with which construct a Markov state model. **A)** Definition of states in the ELViM projection with MSM equilibrium probabilities after Maximum Caliber reweighting with a schematic representation of the domain rearrangements of GSN domains per state. B) Schematic picture of the reweighted MSM using the 75 percentiles of the most probable transition (for visualization purposes) with a schematic representation of the domain rearrangements of GSN domains per state. **C)** Fraction of domain G2-G3 (G_23_) and G4-G6 (G_456_) RMSD (RMSD≤0.55nm) from the actin-capping (actin-bound) or closed conformations. For each state, the first column represents G_23_ and the second G_456._ The fraction of the domains is used to define the schematic state in panels A and B.

This structural interpretation is consistent with DEER measurements. Distances probing the N-terminal block (Nter-G2, Nter-G3, and G1-G3) display broad and heterogeneous distributions, whereas distances reporting on the G4-G6 region (G4-G5 and G4-C_ter_) indicate a more restricted conformational variability. To further investigate the dynamics of the ensemble, we constructed a Markov State Model (See Methods 12) using the deeptime python library (52). To account for the ensemble weights, we used a Maximum Caliber (53) approach by reweighting the inferred transition rate matrix (54). We analyzed the slowest dynamical modes revealing two dominant motions (Figure S18). The slowest process corresponds to rearrangements within the G4-G6 block, whereas the second slowest process involves relative motions between domains G1 and G2. Together, these results indicate that the calcium-bound open state of gelsolin is characterized by extensive conformational heterogeneity, with a highly dynamic N-terminal region coexisting with a comparatively stable C-terminal block.

## Discussion

Gelsolin is involved in a remarkably broad range of physiological and pathological processes (3, 7) and its biology has been extensively investigated over the last four decades. Structural interpretation of these findings has long relied on a limited set of experimental models, primarily represented by the crystal structure of the compact Ca²⁺-free protein and by structures of isolated Ca²⁺-bound domains or domain triplets (34, 36–39). Only recently, the cryo-EM structure of the F-actin:gelsolin capping complex provided a structural view of an alternative functional state of the protein (40). As a consequence, many previous observations on gelsolin function and regulation can now be revisited in the light of a more complete structural framework. The present study contributes to this effort by providing experimental and computational insights into the activation pathway of full-length gelsolin and into the conformational ensemble populated by this active state.

A mechanistic model of gelsolin activation had previously been proposed, derived mainly from crystallographic structures of the inactive protein and isolated Ca²⁺-bound fragments (2) and therefore focused on local conformational rearrangements and the release of domain-domain contacts and latches that maintain the compact conformation. Our results largely support these models but extend them by providing both a temporal and a global perspective of the activation process. The first major activation event identified by both DEER and simulations is the separation of the two major domain blocks, G1-G2-G3 and G4-G5-G6. While the temporal information provided by simulations cannot be directly mapped onto the Ca²⁺ titration, this transition appears both as the earliest event along the simulated opening pathway and as the first major conformational rearrangement detected upon increasing Ca²⁺ concentration. This process is followed by the appearance of intermediate conformations characterized by partial inter-triplet separation, whereas subsequent rearrangements primarily involve the N-terminal triplet, whose domains progressively dissociate and explore a broad conformational space. In contrast, the G4-G5-G6 block remains largely compact throughout activation, with the notable exception of the highly dynamic C-terminal tail. Together, these observations reveal that gelsolin activation is not a concerted transition but rather a hierarchical process involving successive structural layers of organization. This hierarchy also accounts for the long-standing disagreement about the number of Ca²⁺-induced transitions and the concentrations required to trigger them (28–30, 32): observables sensitive to global size saturate once inter-triplet separation is complete, around 200 µM, whereas local reporters continue to change up to millimolar Ca²⁺, so that the apparent number of transitions depends on the probe rather than on the protein.

Although a systematic characterization of the affinities of the individual calcium-binding sites is still lacking, previous studies suggest that calcium binding may occur according to an affinity hierarchy, with G6 displaying the strongest affinity, followed by G2 and then G4 (summarized in(2)). Within this framework, the concentration dependence observed by DEER suggests a possible interpretation of the activation pathway. The earliest detectable change, involving increased mobility of the C-terminal region reported by the G4-Cter reporter at approximately 100 μM Ca²⁺, may reflect structural consequences of occupancy of the highest-affinity sites. At approximately 200 μM Ca²⁺, where the SAXS signal reaches a plateau, additional changes become apparent in the G3-G5 and G2-Cter reporters, consistent with separation of the two domain triplets and the stabilization of an intermediate state. Complete activation is reached only at millimolar calcium concentrations, suggesting that the full process requires the cumulative effect of multiple binding events.

The calcium dependence measured here can also be read against the functional and the cellular calcium scales. The appearance of partially separated G3–G5 conformers between 0.2 and 2 µM Ca²⁺ coincides with the concentration range in which full-length gelsolin binds F-actin, exposes regions accessible to limited proteolysis and acquires a first actin-binding site (30, 33) and with release of the C-terminal tail latch (32), whereas the half-transition of global opening (K_1/2_ ≈ 18 µM free Ca²⁺) matches the ∼10 µM half-maximal severing and nucleation measured at neutral pH (31). The structural titration therefore reproduces the functional titration: partial inter-triplet separation is sufficient for filament binding, while efficient severing requires the more extensive rearrangements of the N-terminal triplet that are completed only at higher Ca²⁺. Placed in a cellular context, these numbers imply that the two isoforms occupy different regions of the same landscape. Plasma gelsolin, synthesized in the secretory pathway and circulating at ∼1.2 mM ionized Ca²⁺, resides permanently in the fully activated ensemble described here, which therefore represents the resting state of the circulating actin-scavenger protein (17, 18). Cytoplasmic gelsolin, by contrast, rarely experiences more than ∼1 µM Ca²⁺ globally (15) and should populate mainly the partially separated intermediates, with fuller opening expected only within calcium microdomains close to open channels, where the cortical actin remodelling to which gelsolin contributes takes place. The observation that actin lowers the Ca²⁺ requirement for gelsolin activation (30) fits naturally into this picture: filaments can capture sparsely populated open conformers and shift the equilibrium, so that the effective threshold in the cytosol is set jointly by Ca²⁺ and by actin availability, a form of conformational selection (55). We note that K_1/2_ was measured in a defined buffer without Mg²⁺ competition or macromolecular crowding, and that the cellular values above are free concentrations with substantial spread between probes and sub-compartments; the comparison should therefore be read in terms of ranges rather than exact values.

So far, active gelsolin has only been represented through static structural snapshots (20, 35, 38, 39), although these same studies and others on a homologous protein (56) hinted at a progressive loss of interdomain contacts. One of the strongest messages that emerged from this study is that the active state of gelsolin cannot be adequately described by a single structure. Rather than converging toward a unique open conformation, activated gelsolin populates a broad ensemble of states characterized by substantial conformational heterogeneity. This behavior is particularly evident from the Markov state model analysis of the trajectories, which reveals a large range of accessible conformations. The recently reported cryo-EM structure of the F-actin:gelsolin capping complex (40) therefore likely represents one functionally relevant member of a much broader conformational landscape. Our data indicate that this structure should not be interpreted as the sole active state of gelsolin, but rather as one of the conformations accessible to the activated protein.

The existence of such an ensemble raises an important question: why should gelsolin remain highly plastic after activation? Our approach does not assay domain local flexibility, but previous studies have never evidenced increased dynamics at the level of the individual modules, rather the opposite: calcium ions have a structural role for the isolated G2 domain, for example (57, 58). Conformational diversity originates primarily from the relative arrangement of the rigid modules connected by mobile linkers. In such a dynamic landscape, some conformations are substantially more populated than others. In particular, the states identified here as S8 and S12 together account for approximately 40% of the equilibrium ensemble. These predominant states are characterized by a C-terminal half that stably adopts a conformation closely resembling the actin-capping structure, while the N-terminal domains are dissociated from one another and maintain broad exposure of their actin-binding surfaces. Neither the closed conformation nor that observed in the capping complex are significantly populated, nor are they directly detected by the DEER reporters, suggesting that Ca^2+^ activation, inactivation by inositol phosphates or ATP, and F-actin binding, whether for capping or severing, all proceed by selection and stabilization of pre-existing sub-states rather than by switching between two discrete conformations (55). In this regard it is notable that the phosphoinositide-binding sequences of gelsolin, residues 135– 142 and 150–169 (21, 59), lie precisely in the G1–G2 region that our data identify as the most plastic part of the activated ensemble: the same mobility that allows the N-terminal triplet to search for productive filament geometries may be what makes it available for compaction by phosphoinositides even at saturating Ca²⁺ (21–23). A single plastic landscape, biased in different directions by Ca²⁺, lipids, nucleotides, pH and proteolysis, offers a more economical description of gelsolin regulation than a set of independent switches.

Such high plasticity may also represent an evolutionary solution to the diverse structural contexts in which gelsolin operates. Gelsolin must not only perform both severing and capping activities but must do so in the presence of competing actin-binding proteins and within cytoskeletal architectures that can vary dramatically between cell types and physiological conditions. Indeed, recent reviews and editorials (60, 61) report 160 actin-binding proteins, classified based on their function including promoting nucleotide exchange and phosphate dissociation, initiating polymerization, capping filament ends and cross-linking filaments to each other or to other proteins. Moreover, actin filaments are organized into bundles (62), branched networks (63), stress fibers (64) and other higher-order assemblies, each imposing distinct steric and geometric constraints. A structurally plastic active state may therefore maximize the probability of productive binding across a wide range of actin environments.

Our results further suggest that different gelsolin domains play distinct mechanistic roles during activation and interaction with actin. The combined analysis of the calcium titration, the simulated opening sequence, and the conformational dynamics reveals a clear functional asymmetry within the protein. This view is consistent with recent cryo-EM studies showing that G2-G3 act as an F-actin recognition module and prime the filament for severing (41). Our data extend this model by suggesting that G2 exposes its actin-binding surface early during activation and that it remains largely accessible at equilibrium, consistent with a pioneer role. The G4-G6 triplet undergoes relatively limited internal rearrangements once activation is complete. This observation agrees well with the available structural and biochemical evidence indicating that this region contributes mainly to capping activity by stabilizing interactions at the filament end and sterically preventing repolymerization (40) but it is not endowed with severing activity (65).

In contrast, the G1-G3 triplet displays substantially greater conformational heterogeneity. Among these domains, G1 appears particularly dynamic and may play a more active role in filament remodeling. Previous structural studies of the capping complex suggested that G1 contributes directly to filament deformation (40), and our observations are consistent with this hypothesis. The marked mobility of the G1-G3 region may be required to adopt the multiple geometries necessary for productive engagement of actin filaments during severing. G1 can be considered the catalytic module. Removal of G1 abolishes severing activity, indicating that the N-terminal triplet is essential for function, but its activity is partially restored when it is replaced by G4 (41). This observation suggests that severing activity is not entirely encoded by the specific sequence or structure of G1 itself. Rather, broader biophysical properties, including the enhanced structural plasticity of the triplet revealed by this study, may contribute to function. The paralogue CapG, which comprises three domains homologous to G1–G3 and caps but does not sever actin filaments, supports this view: severing can be conferred on CapG by grafting gelsolin sequences that lengthen the arm connecting its first two domains, and is lost again upon insertion of a single residue into that arm (66), indicating that the geometry and mobility of the G1–G2 linkage, rather than the actin-binding surfaces as such, encode severing competence. The resulting division of labour, a pre-organized C-terminal half that anchors at the barbed end and a conformationally searching N-terminal half that cuts, is reminiscent of the fly-casting mechanism (67), in which a flexible module enlarges the capture radius for a partner while a rigid module provides a binding site with a low entropic cost.

These findings may also help reinterpret several pathological processes involving gelsolin. A notable example is AGel amyloidosis (reviewed in (68)), a disease extensively studied by some authors of the present work. In this condition, pathogenic substitutions in G2 allow furin to cleave plasma gelsolin at Arg172 during transit through the trans-Golgi network, because the mutant domain is no longer stabilized by Ca²⁺ at the concentration prevailing in that compartment (57, 69). Our data add the global counterpart to this local-stability argument: at the ∼100–300 µM free Ca²⁺ of the Golgi lumen (16), wild-type gelsolin is predicted to have completed inter-triplet separation but not the full rearrangement of the N-terminal triplet, so that the species presented to furin is a specific, partially open intermediate in which G2 is already released from the C-terminal half and its accessibility is governed by its own Ca²⁺-dependent stability. The same framework predicts that substitutions weakening G2 Ca²⁺ binding should shift K_1/2_ and the population of this intermediate, a testable consequence. More generally, the lack of structural information on activated gelsolin has likely limited efforts to exploit the protein as a therapeutic target, whether by stabilizing selected states of the cytoplasmic isoform or by engineering the recombinant plasma protein now in clinical development (18). By identifying domain-specific activation mechanisms and the conformational states populated along the activation pathway, the present work provides a structural framework that may facilitate such strategies.

Despite the advances presented here, critical structural information is still missing. In particular, the molecular architecture of the severing complex remains unknown. Such a structure would provide the ultimate test for the functional hypotheses proposed for the different gelsolin domains and would clarify how the conformational plasticity of the active state is translated into actin filament remodeling.

## Supporting information

supplementary results and methods

## Acknowledgment

We acknowledge the European Synchrotron Radiation Facility (ESRF) for provision of synchrotron radiation facilities under proposal MX-2691 and MX-2591. We thank the scientist of beamline BM29 for assistance and support during the beamtime.

The authors would also like to thank Diamond Light Source for beamtime (proposal MX20221) and the staff of beamlines B21 for assistance with data collection.

The authors acknowledge CINECA for an award under the ISCRA initiative, for the availability of high-performance computing resources and support.

The EPR facilities at the French Research Infrastructure INFRANALYTICS(FR2045) and the Aix-Marseille University EPR Center are also acknowledged by the authors.

This project has received funding from the European Union’s Horizon 2020 research and innovation programme under grant agreement No 101004806. We acknowledge TNA to the MOSBRI EPR infrastructure under project n° MOSBRI-2023-168.

This work was supported by the National Recovery and Resilience Plan (NRRP), Mission 4, Component 2, Investment 1.1, Call for tender No. 104 published on 2.2.2022 by the Italian Ministry of University and Research (MUR), funded by the European Union – NextGenerationEU – Project Title “Novel Approaches to Gelsolin Amyloidosis Treatment – NAGAT” – CUP B53D23015890006;

This work was supported by Fondazione Telethon ETS-Italy (Grant no. GMR24T1097).

## Author contribution

BS, LG, AB, CC, MdR designed research; BS, LG, FB, GG, MB, AB performed research; BS, LG, TG, RC, MM, AB, CC, MdR analyzed data; BS, MM, AB, CC, MdR wrote the paper; all authors edited and revised the paper

