## supplementary results and methods for "A rigid anchor and a flexible blade: calcium activates gelsolin through an asymmetric conformational ensemble"

| | |
|---|---|
| **Supplementary results** | **pg 2** |
| Supplementary SAXS data (Table S1, Figure S1) | pg 2 |
| Supplementary DEER data (Figure S2-S3) | pg 4 |
| Multi-eGO model and preparation (Figure S4-S5) | pg 6 |
| Model Validation | |
| SAXS BME (Figure S6-S9) | pg 7 |
| DEER (Figure S10-S12) | pg 9 |
| Solvent Accessible Surface Area (SASA, Figure S13-14) | pg 14 |
| Energy Landscape Visualization Method (ELVIM, Fig S15-S25) | pg 16 |
| SAXS data fitting (Fig S27, Table S2) | pg 24 |
| **Supplementary methods** | **pg 25** |
| **Supplementary References** | **pg 37** |

### Supplementary results

#### Supplementary SAXS data

***Table S1:*** *Global structural parameter derived from SAXS data by varying GSN and $Ca^{2+}$ concentration. The radius of gyration ($R_g$) and forward scattering intensity ($I_0$) were calculated from the Guinier approximation, Dmax from the pair-distance distribution function. Data processing was performed with PRIMUS (ATSAS software suite, [1])*

| $Ca^{2+}$ ($\mu$M) | GSN ($\mu$M) | $Ca^{2+}$/GSN molar ratio | $R_g$ (Å) | I0 (a.u.) | Dmax (Å) | SASDB id |
|---|---|---|---|---|---|---|
| 0 | 46.25 | 0 | 31.5±0.2 | 0.058 | 93.9 | SASD2Z8 |
| 0 | 23.54 | 0 | 31.8±0.2 | 0.057 | 103.9 | |
| 0 | 10.77 | 0 | 31.5±0.2 | 0.056 | 103.8 | |
| 0 | 5.27 | 0 | 31.9±0.2 | 0.059 | 101.5 | |
| 0 | 2.69 | 0 | 31.7±0.3 | 0.054 | 99.1 | |
| 50 | 52.11 | 1 | 32.7±0.7 | 0.050 | 97.0 | SASD269 |
| 50 | 27.52 | 2 | 33.9±0.3 | 0.055 | 116.7 | |
| 50 | 16.63 | 3 | 34.8±1.0 | 0.050 | 105.0 | |
| 50 | 8.43 | 6 | 36.9±0.2 | 0.045 | 111.6 | |
| 50 | 4.45 | 11 | 40.0±0.5 | 0.034 | 117.7 | |
| 200 | 48.83 | 4 | 41.0±0.5 | 0.066 | 144.4 | SASD2B9 |
| 200 | 25.76 | 8 | 41.2±0.5 | 0.062 | 140.3 | |
| 200 | 14.17 | 14 | 40.7±0.3 | 0.057 | 124.0 | |
| 200 | 7.61 | 27 | 40.1±0.2 | 0.051 | 128.4 | |
| 200 | 3.86 | 52 | 41.4±0.7 | 0.051 | 124.3 | |
| 1000 | 48.13 | 21 | 42.2±0.5 | 0.065 | 136.5 | SASD2G9 |
| 1000 | 25.88 | 39 | 41.3±0.5 | 0.060 | 140.6 | |
| 1000 | 13.58 | 74 | 40.8±0.3 | 0.059 | 127.8 | |
| 1000 | 7.03 | 142 | 39.8±0.3 | 0.055 | 127.3 | |
| 1000 | 3.51 | 285 | 39.3±0.5 | 0.050 | 117.1 | |
| 10000 | 46.84 | 214 | 41.9±0.4 | 0.063 | 131.6 | SASD2M9 |
| 10000 | 24.47 | 409 | 41.0±0.7 | 0.059 | 130.4 | |
| 10000 | 13.23 | 756 | 40.3±0.5 | 0.055 | 123.8 | |
| 10000 | 6.91 | 1447 | 39.8±0.3 | 0.051 | 130.7 | |
| 10000 | 3.86 | 2588 | 41.2±0.4 | 0.046 | 123.1 | |

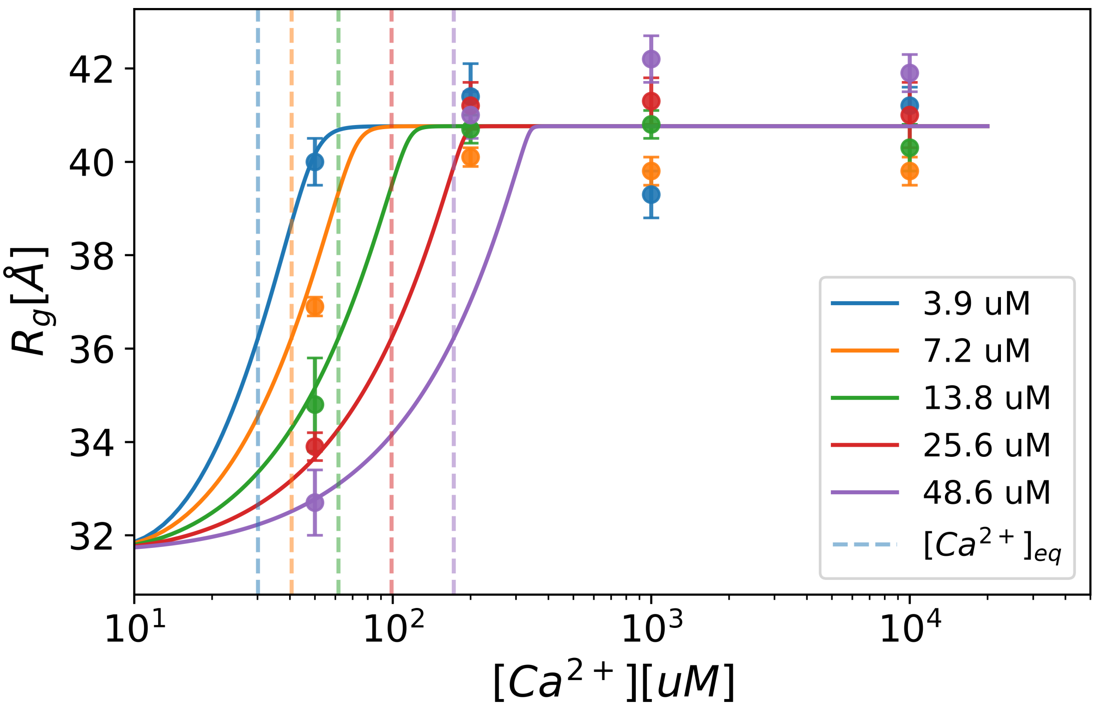


***Figure S1:*** *All $R_g$ Vs. $[Ca^{2+}]_{tot}$ data reported in Table S1 are compared with the curves predicted by the two-state model described in the main text, using $K_{1/2}$ = 18 ± 4 μM and $n_{eff}$ = 6.4 ± 0.9 parameters, obtained from the fit of the dataset collected at fixed $Ca^{2+}$ = 50 μM (Figure 1A).*

#### Supplementary DEER data

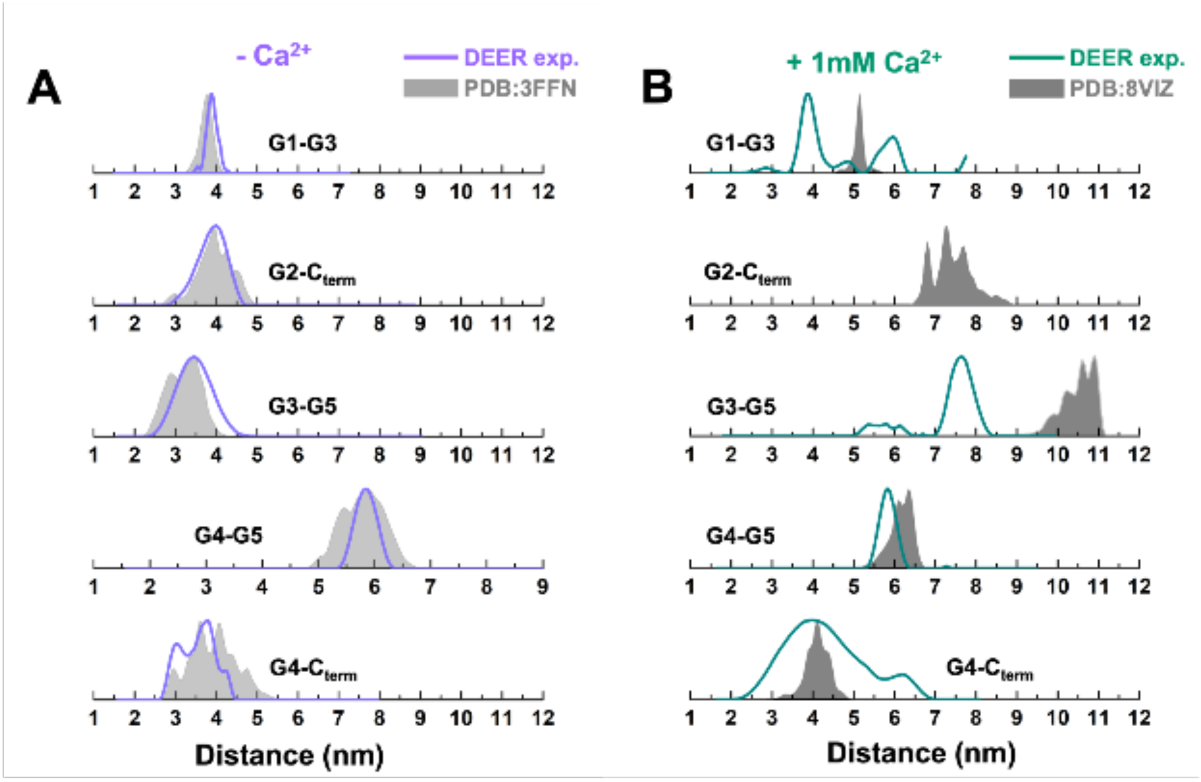


***Figure S2:*** *Selected DEER distances (violet and green) compared to spin-spin distribution computed with MMM2019 (gray) obtained respectively for the closed **(A)** and the actin capping **(B)** states of GSN. In-silico distance distributions have been computed from the structural model available of the full-length system for its closed (PDB: 3FFN) and actin-capping (PDB: 8VIZ) conformations.*

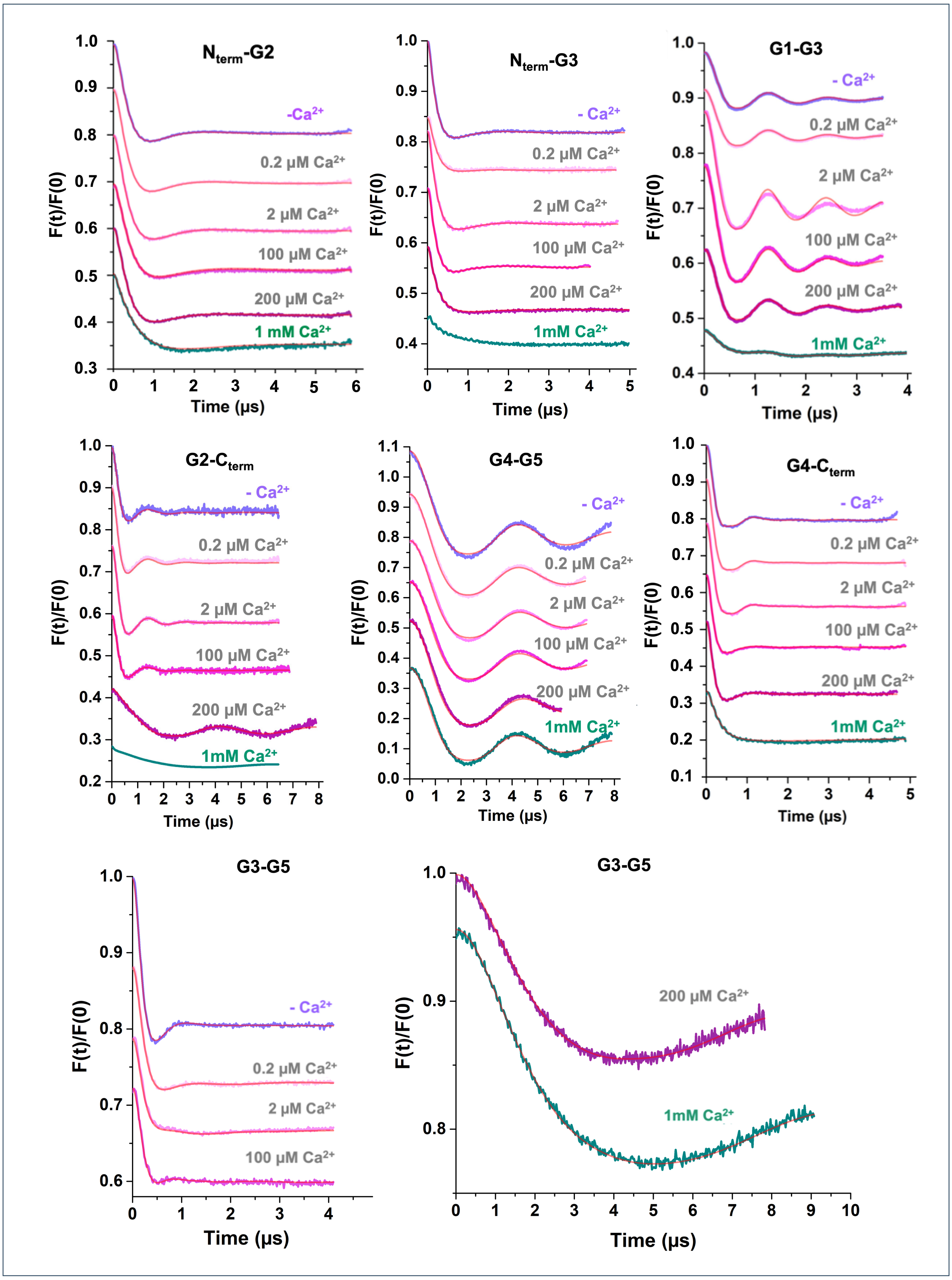


***Figure S3:*** *Background corrected Q-band DEER traces and related fitting curves (red) in the absence and in the presence of increasing concentration of* $Ca^{2+}$ *obtained for the spin labelled GSN mutant bearing the tags respectively in the domain reported above each panel. Lower DEER modulation depth, i.e. a reduction in the amplitude of the DEER oscillations, may arise from very long interspin distances or from highly heterogeneous conformational ensembles.*

### Multi-eGO preparation

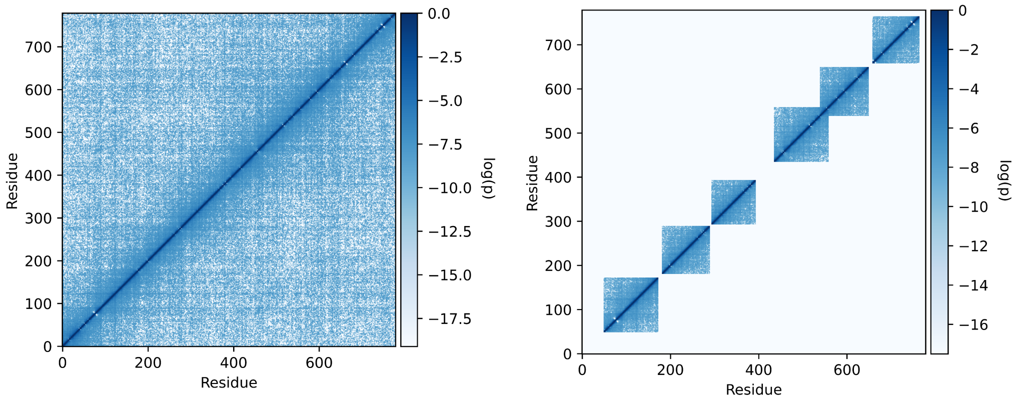


***Figure S4:*** *Matrix representation of Full-length molten globule prior simulation. Intra-domain selection of full-length prior matrix used to define and reweight intra-domain interactions in the inter-domain and production force-field generation.*

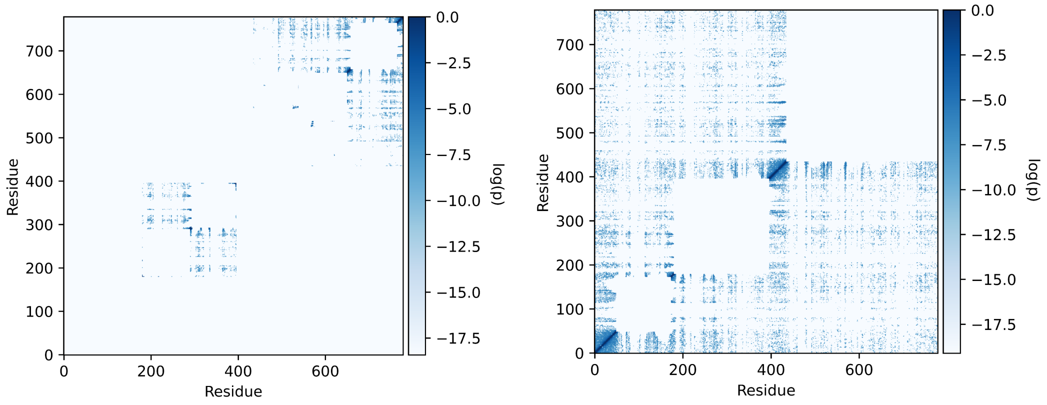


***Figure S5:*** *Inter-domain matrix representation obtained from the inter-domain prior simulations. On the left are shown the contacts between coupled domain interfaces (G2-G3 and G4-G5-G6) and on the right the contacts between all the other domains.*

### SAXS BME

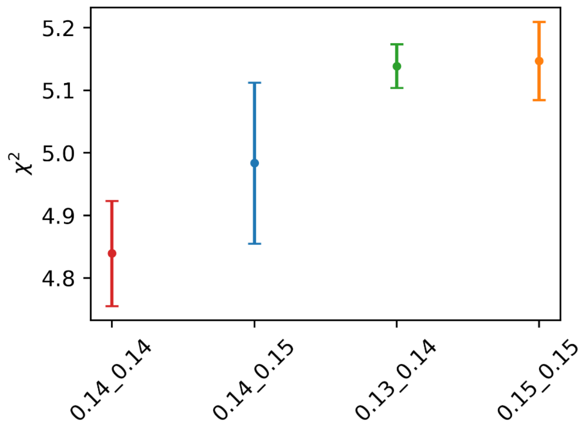


***Figure S6:*** *Unbiased SAXS $\chi^2$ of the different tested set-ups.*

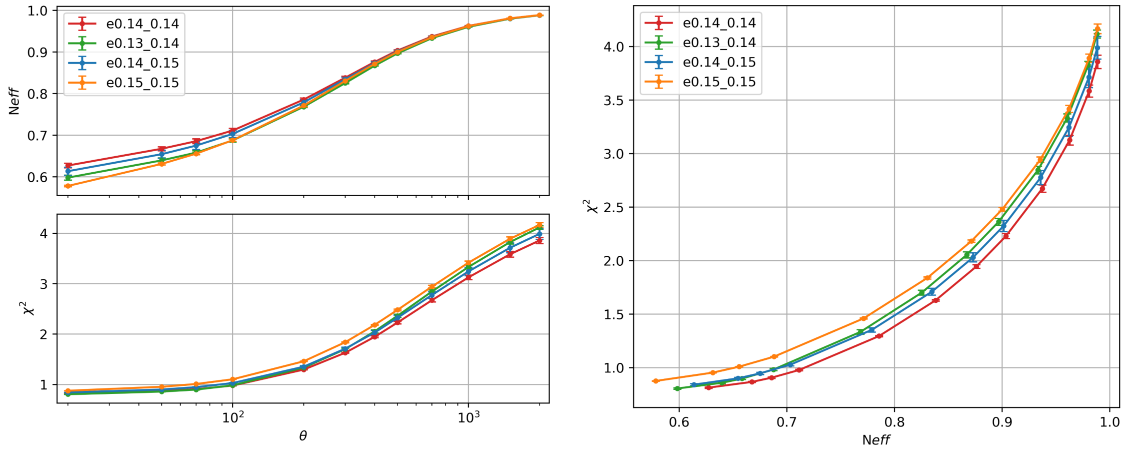


***Figure S7:*** *Agreement between in silico and experimental SAXS profile as a function of the BME scaling parameter θ. On the left are shown the effective number of frames and the $\chi^2$ as a function of θ. On the right is shown instead the $\chi^2$ as a function of the effective number of frames $N_{eff}$. An agreement of $\chi^2 \simeq 1$ is obtained for $\theta \simeq 100$ for all tested set-ups. The set up a) corresponding to 0.14-0.14 shows slightly better behaviour, maintaining a larger effective number of frames given the same value of θ.*

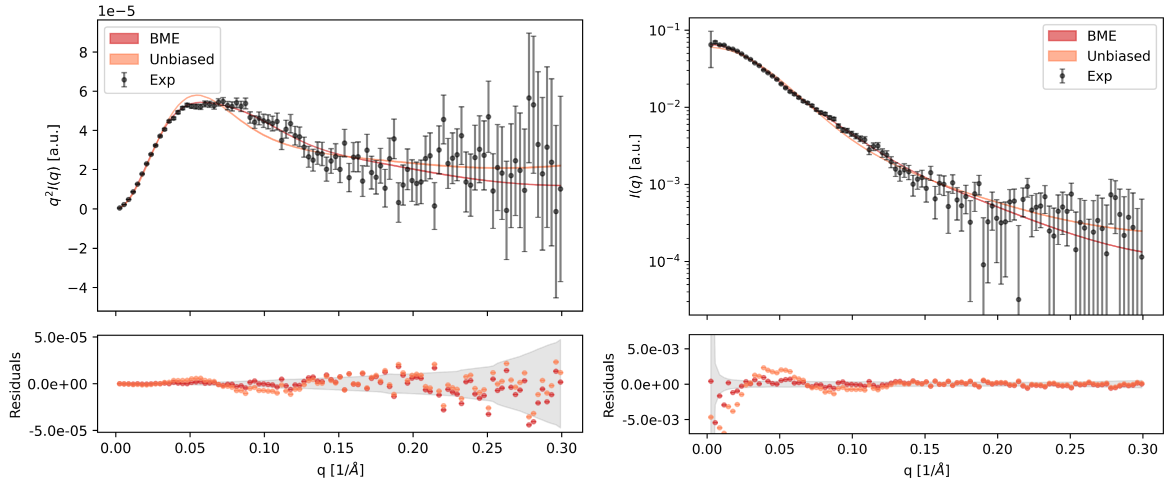


***Figure S8:*** *Experimental and in silico I(q) and $q^2$ I(q) SAXS profile comparisons for set-up a) 0.14-0.14 with and without BME reweighting.*

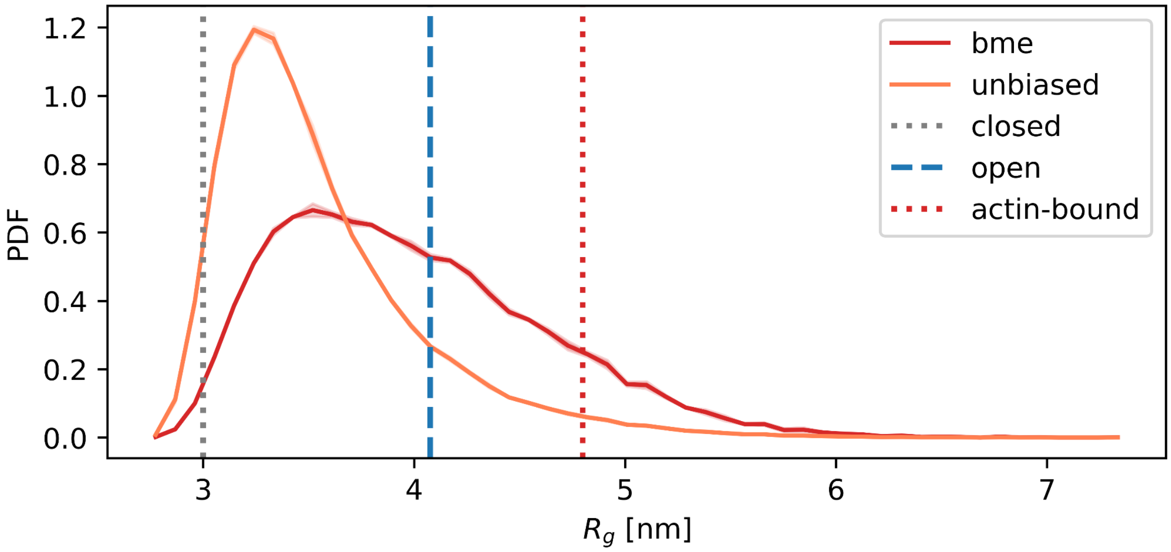


***Figure S9:*** *In silico radius of gyration distribution comparison with and without BME reweighting. Comparison, $R_g$ values of reference structures (closed: PDB: 3FFN, actin-bound PDB: 8VIZ) are shown with dotted vertical lines. Reference average experimental $R_g$ of GSN in the open state (saturation concentration of ions 1mM) is shown with dashed vertical line.*

### DEER

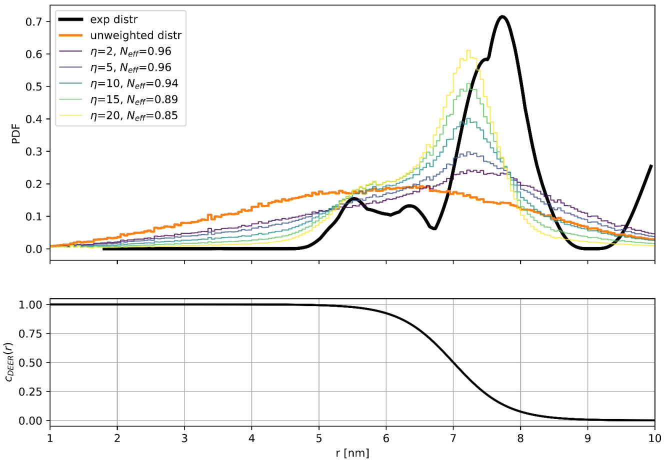


***Figure S10:*** *Comparison between G3-G5 Experimental DEER distance distribution (mean + error) in black line with the in-silico ones calculated using DEERPREdict. In orange is shown the unbiased distribution which populates close distances which are not observed in the experimental distribution. In the other colours are shown the filtered distance profiles as a function of the control parameter* $\eta$ *and the effective number of frames. The bottom panel represents the DEER confidence as a function of distance used in the calculation of the filtering scheme (See Methods).*

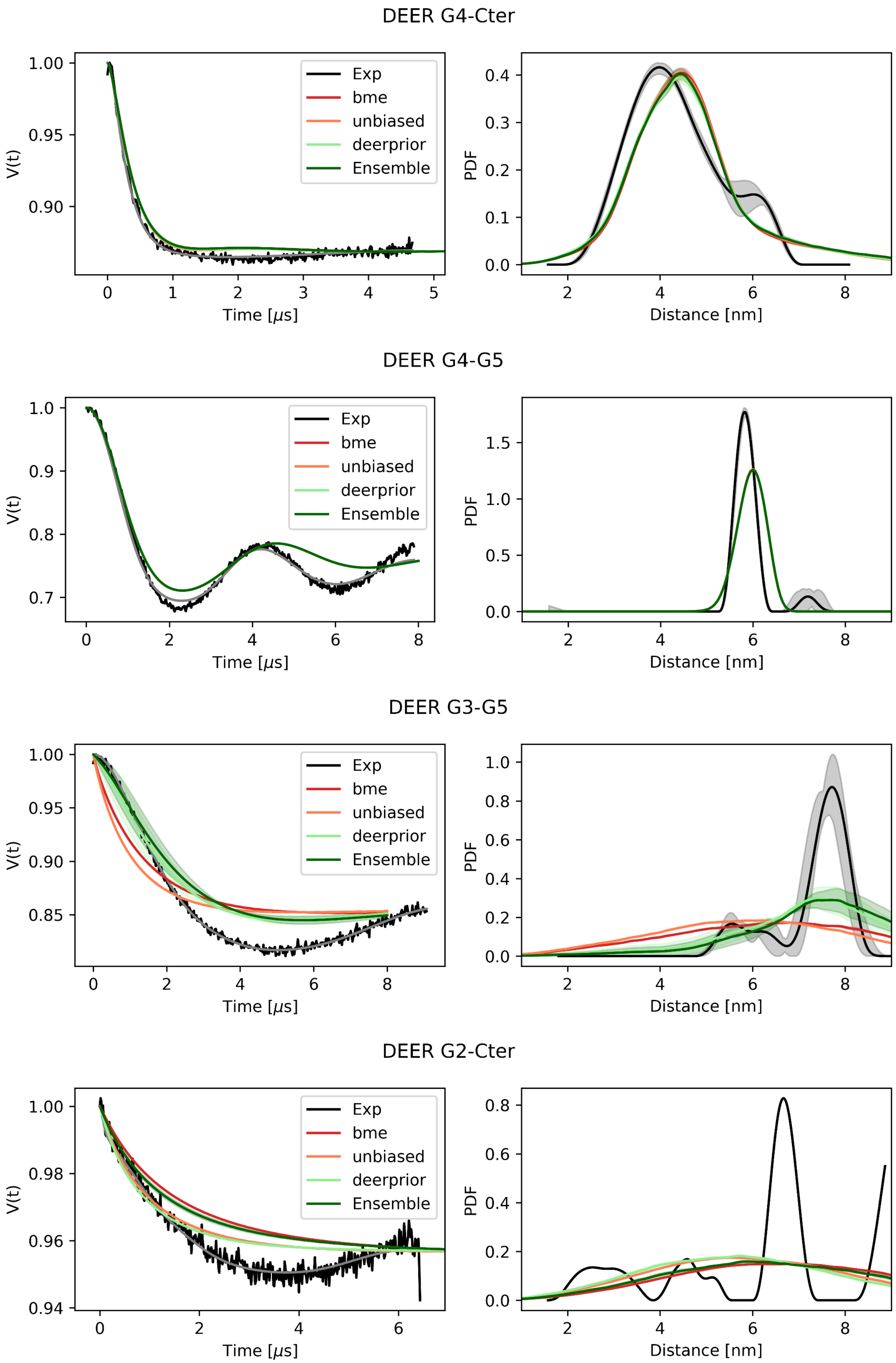
DEER G4-Cter
DEER G4-G5
DEER G3-G5
DEER G2-Cter
Exp
bme
unbiased
deerprior
Ensemble
V(t)
Time [$\mu$s]
PDF
Distance [nm]

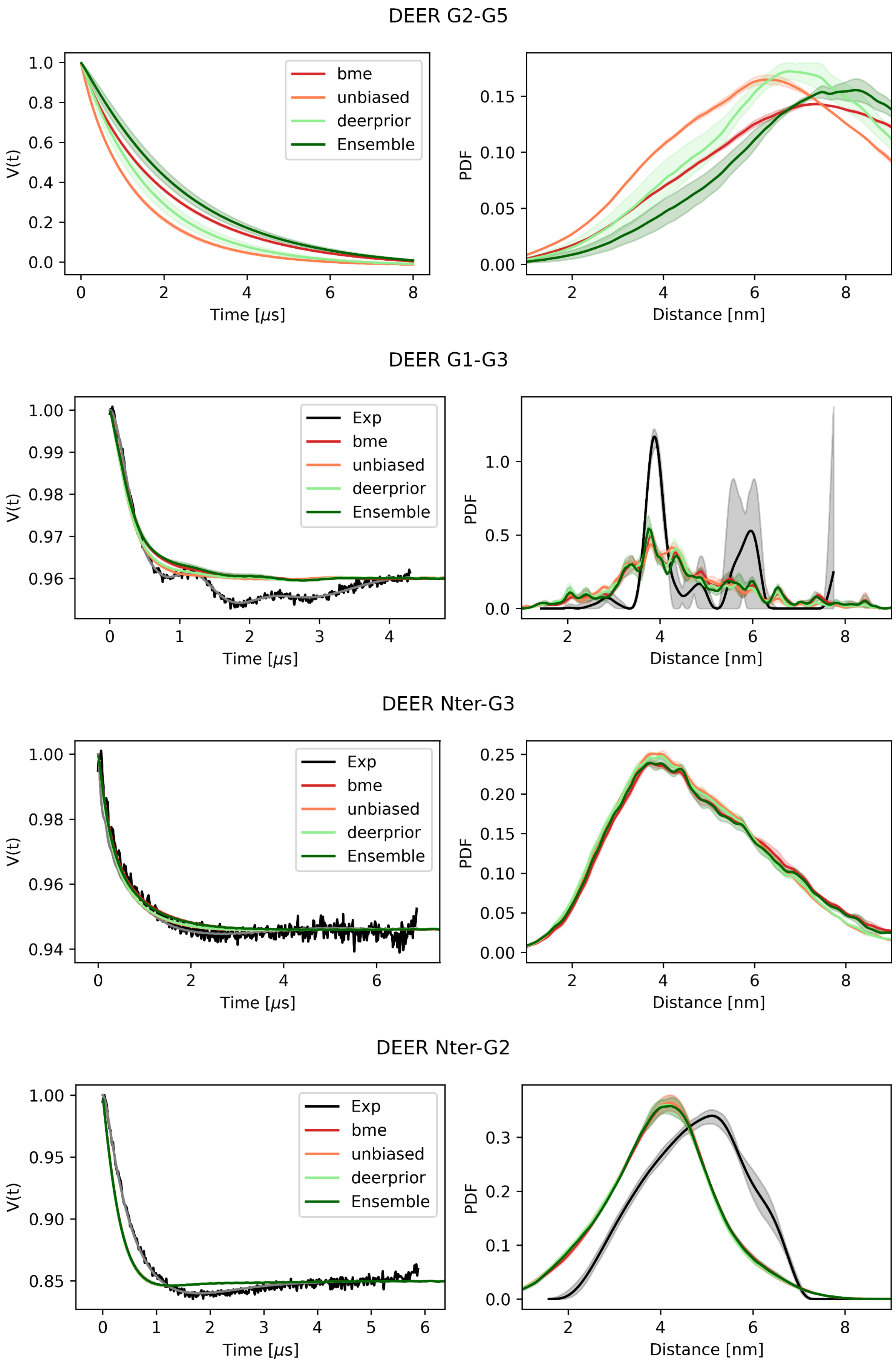
DEER G2-G5
V(t)
Time [μs]
bme
unbiased
deerprior
Ensemble
PDF
Distance [nm]
DEER G1-G3
V(t)
Time [μs]
Exp
bme
unbiased
deerprior
Ensemble
PDF
Distance [nm]
DEER Nter-G3
V(t)
Time [μs]
Exp
bme
unbiased
deerprior
Ensemble
PDF
Distance [nm]
DEER Nter-G2
V(t)
Time [μs]
Exp
bme
unbiased
deerprior
Ensemble
PDF
Distance [nm]

***Figure S11:*** *All DEER profiles comparisons between experimental traces and in silico ones for all the reweighting schemes: Unbiased (no reweighting), bme (SAXS reweighting), deerprior (filtering DEER G3-C5) and Ensemble (combination of prior filtering DEER G4-Cter and SAXS bme reweighting). DEER profiles of G2-G5 have no experimental counterpart.*

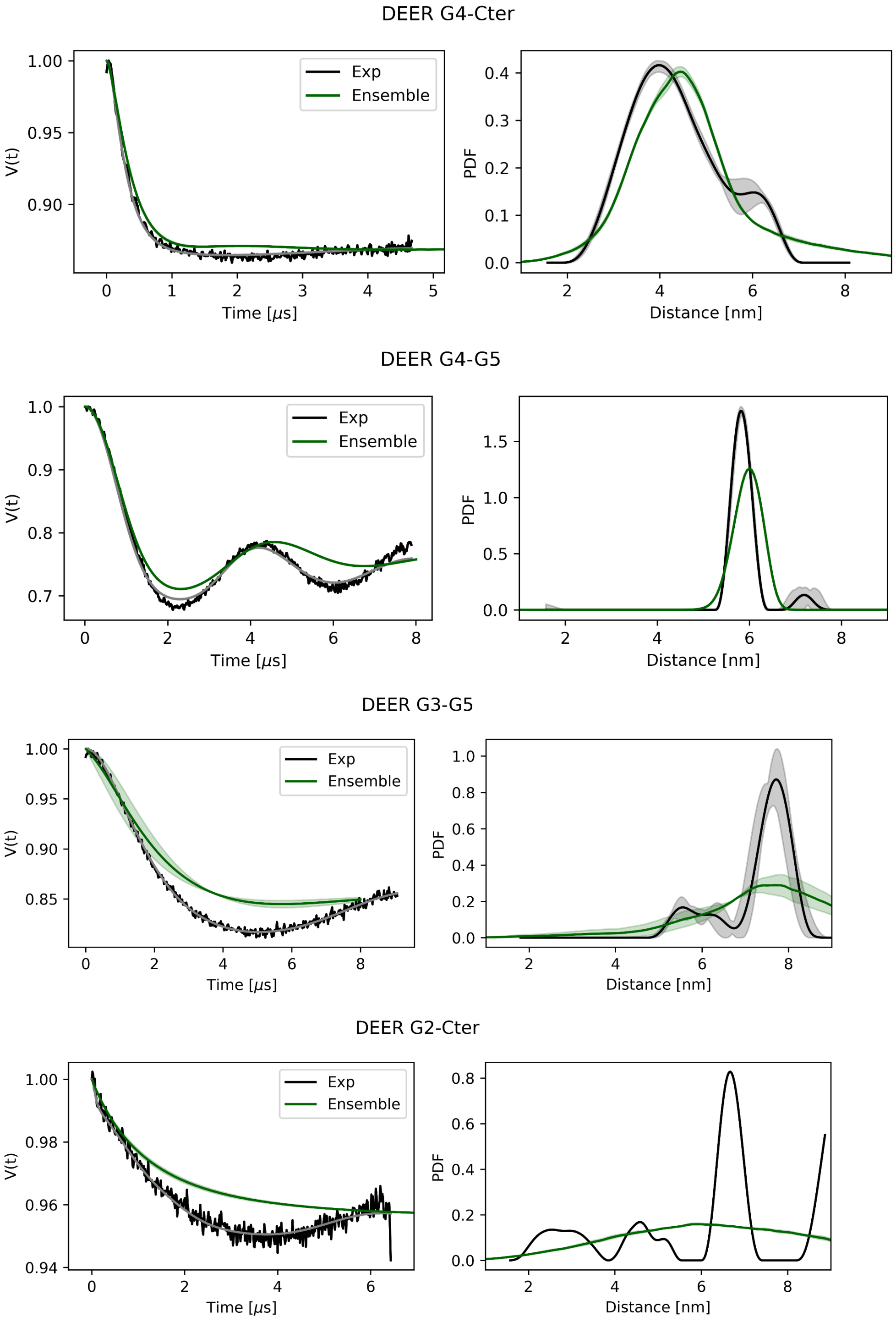

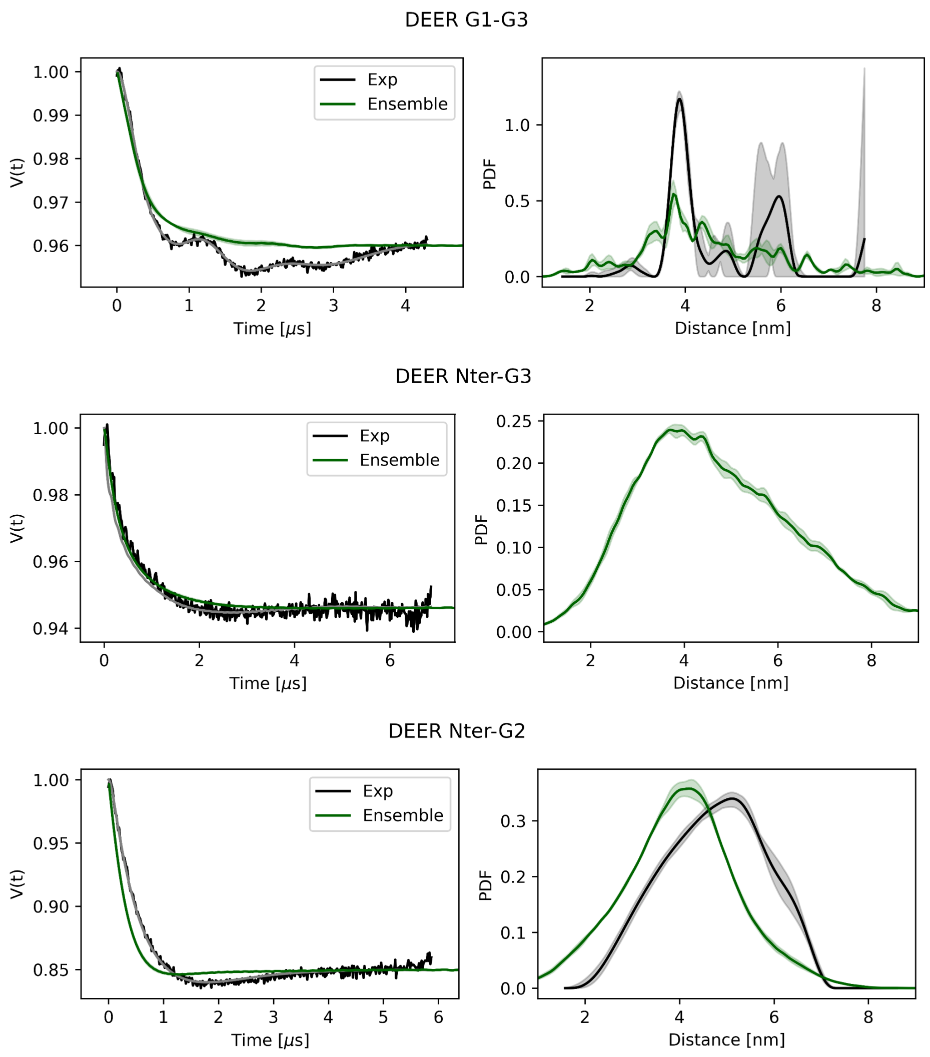


***Figure S12:*** *All DEER profiles comparisons between experimental (black) and in silico (Green) traces of the final Ensemble. DEER profiles of G2-G5 have no experimental counterpart.*

### SASA

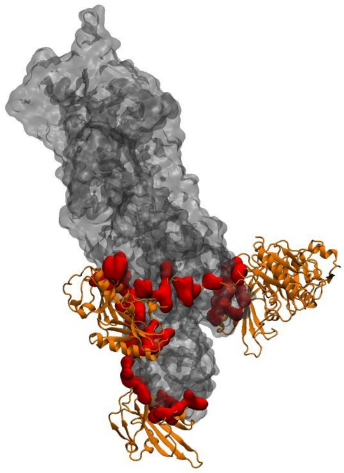

***Figure S13:*** *Visualization of contact residues between GSN and the actin filament in Experimental structure (PDB: 8VIZ) (See Methods)*

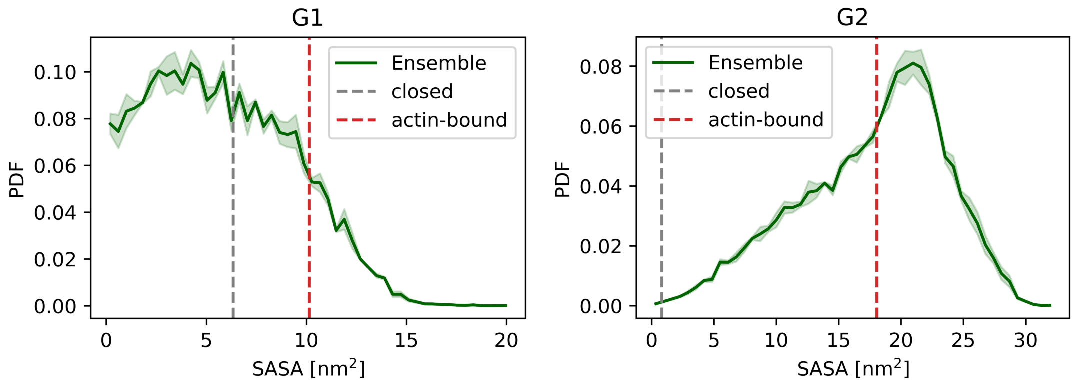

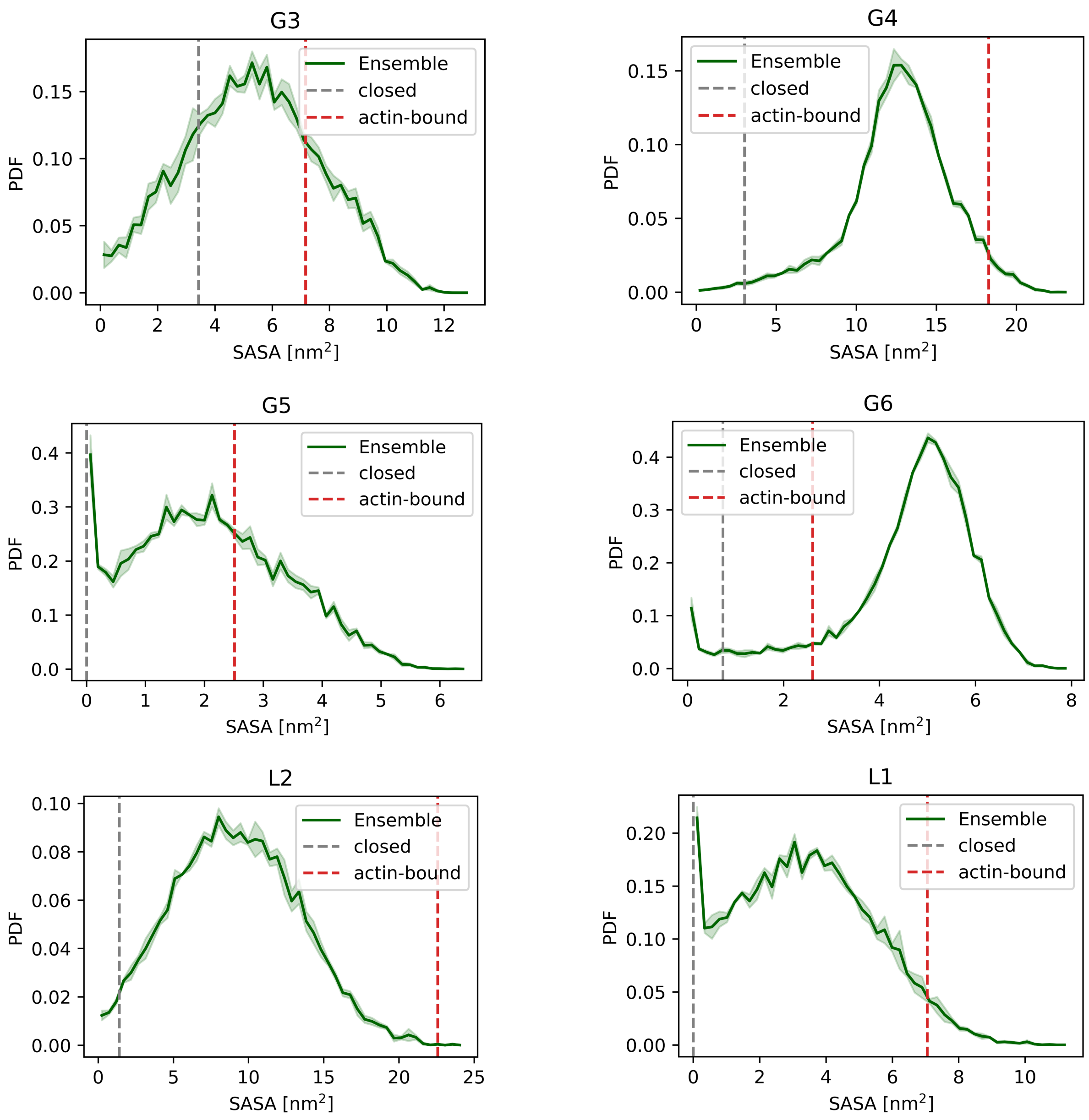


***Figures S14:*** *Solvent Accessible Surface Area (SASA) ensemble distribution for each domain calculated on the residues that are in contact with the actin-filament (See Methods). Comparison values are shown in dashed lines computing the SASA of the same residue from the reference structures of the closed (PDB: 3FFN) and actin bound (PDB: 8VIZ) structures.*

#### ELViM

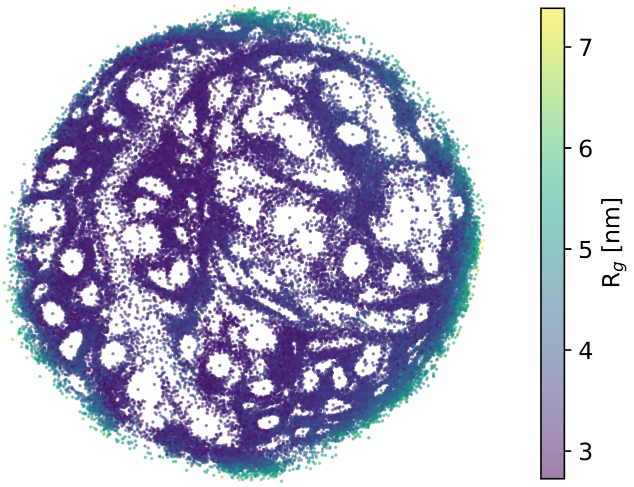


***Figure S15:*** *Radius of gyration mapped on ELViM projection. Expanded conformations lie on the external surface of the projection.*

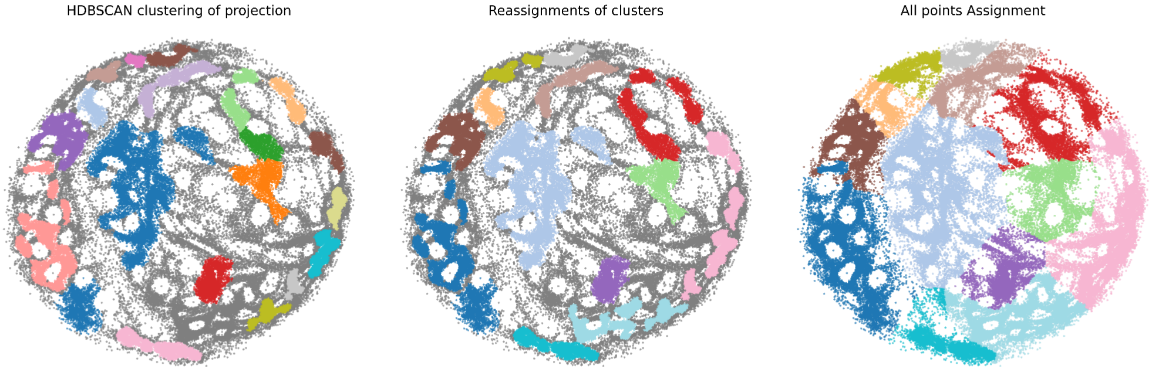


***Figure S16:*** *HDBSCAN clustering steps. a) Starting clusters obtained with HDBSCAN (left) shown in different colours (grey dots is considered noise). b) Merging of clusters (center) based on structural similarities (domain rearrangements and radius of gyration). c) All points assignment (right) used to construct the Markov State Model.*

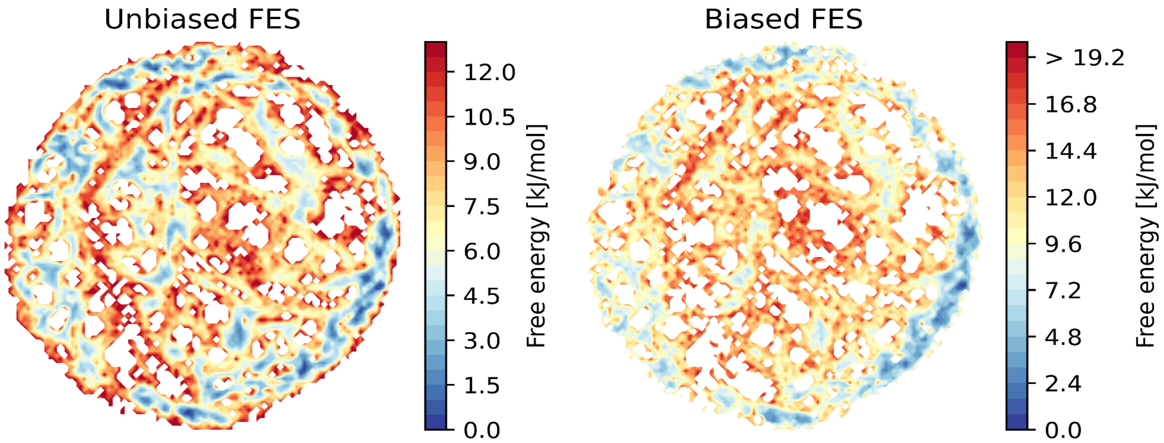


***Figure S17:*** *Free energy surface (FES) of ELViM projection. On the left the unbiased one, and on the right the one reweighted with the final BME DEER+SAXS weights. The overall effect of the reweighting is to disfavor population in the central region of the projection which is associated with more compact structures.*

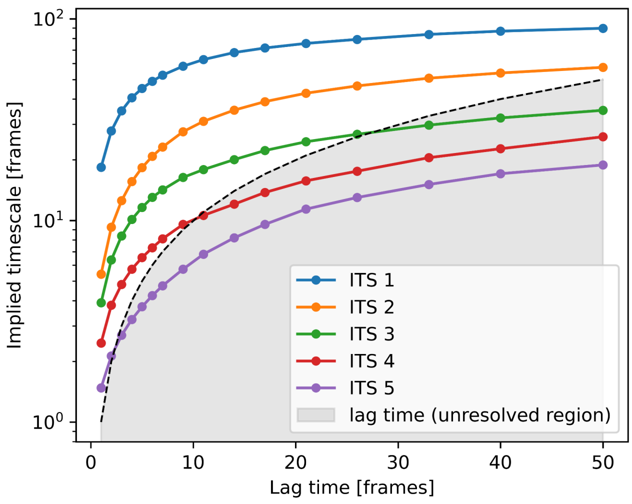


***Figure S18:*** *Implied timescales plot as a function of the lag time. The plot shows 2 clear resolved imlied timescales (IT1, IT2: blue and orange) and a third one that reaches a plateau at lag time 25 (frames) just before the unresolved region (IT3: green)*

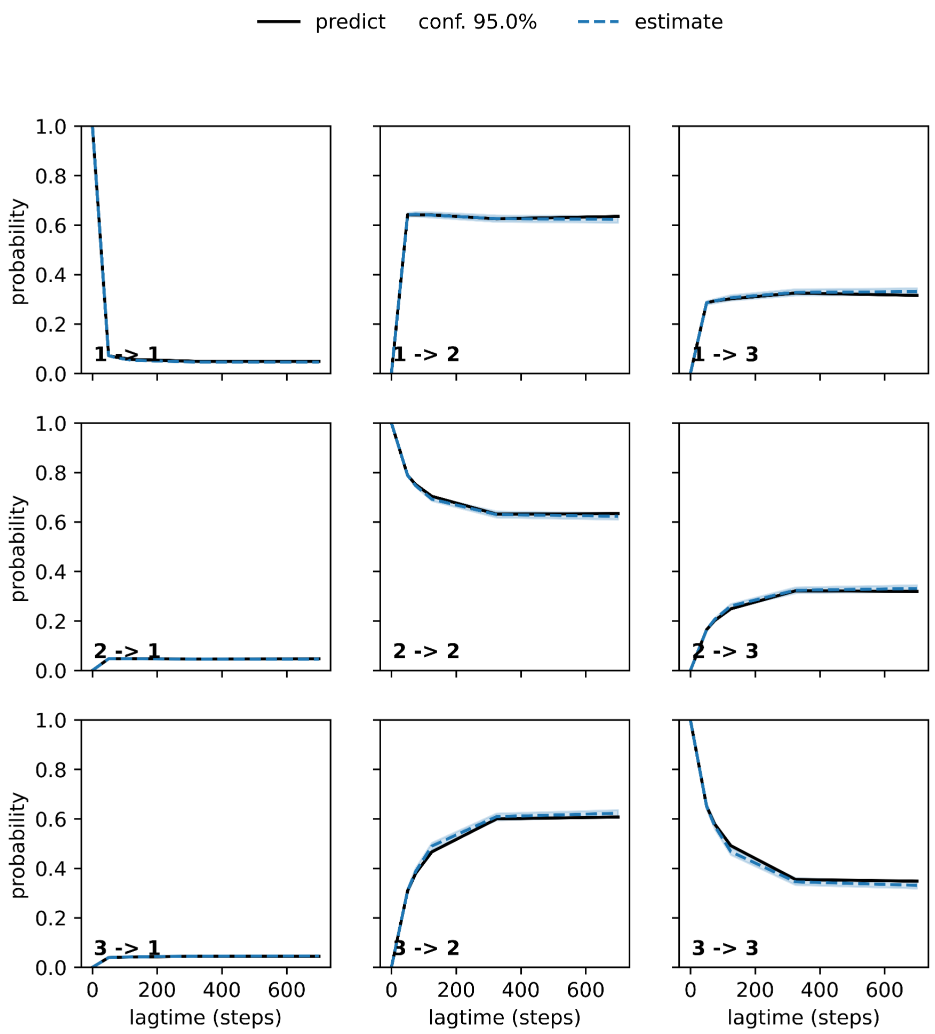


***Figure S19:*** *Chapman-Kolmogorov test of the Markov-State Model using 3 metastable states. The test shows that markov state model correctly describes the time evolution of the system.*

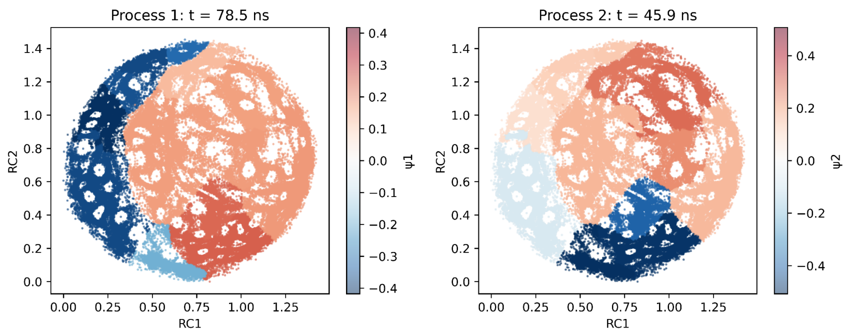


***Figure S20:*** *Eigenvalues projection of the 3 slow implied timescales.*

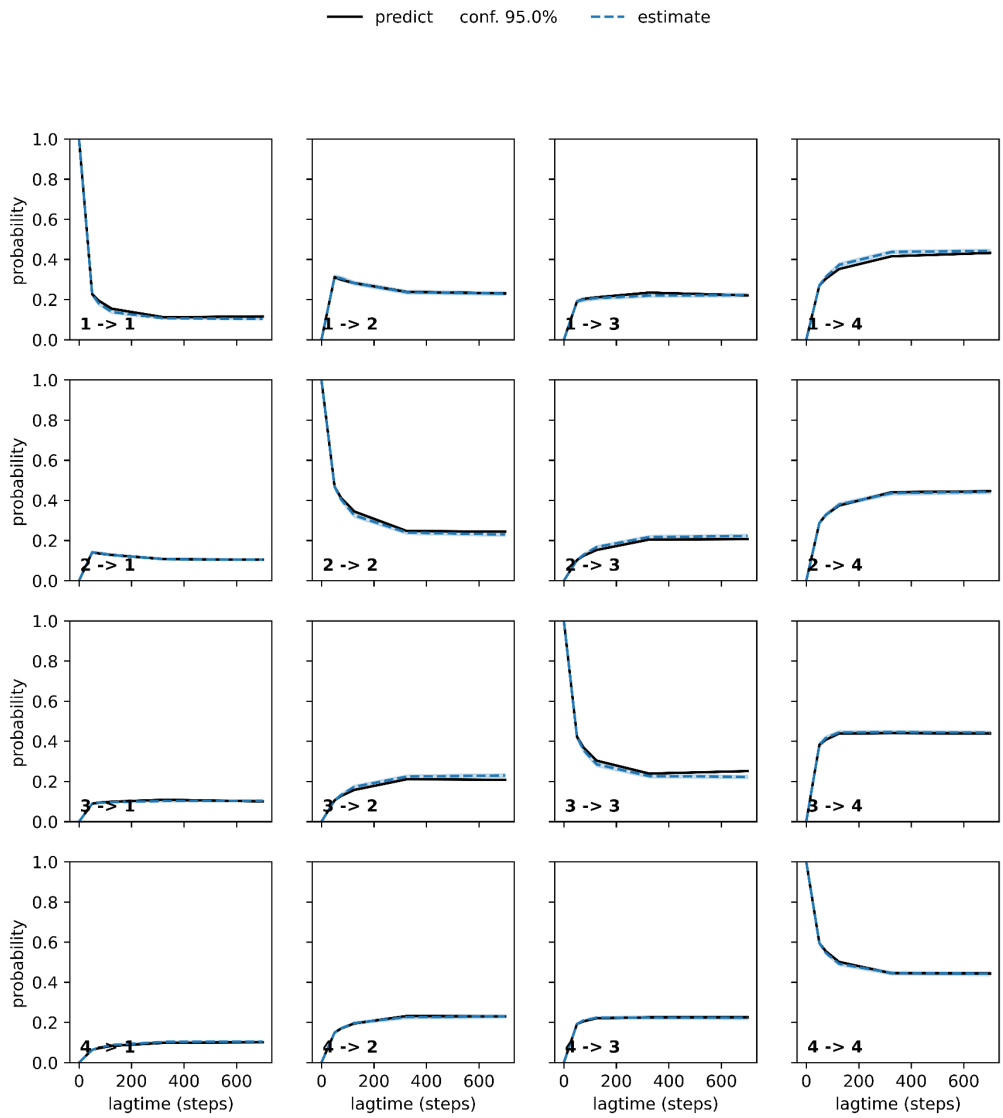


***Figure S21:*** *Chapman-Kolmogorov test of the Markov-State Model using 4 metastable stateds. The test shows that markov state model correctly describes the time evolution of the system.*

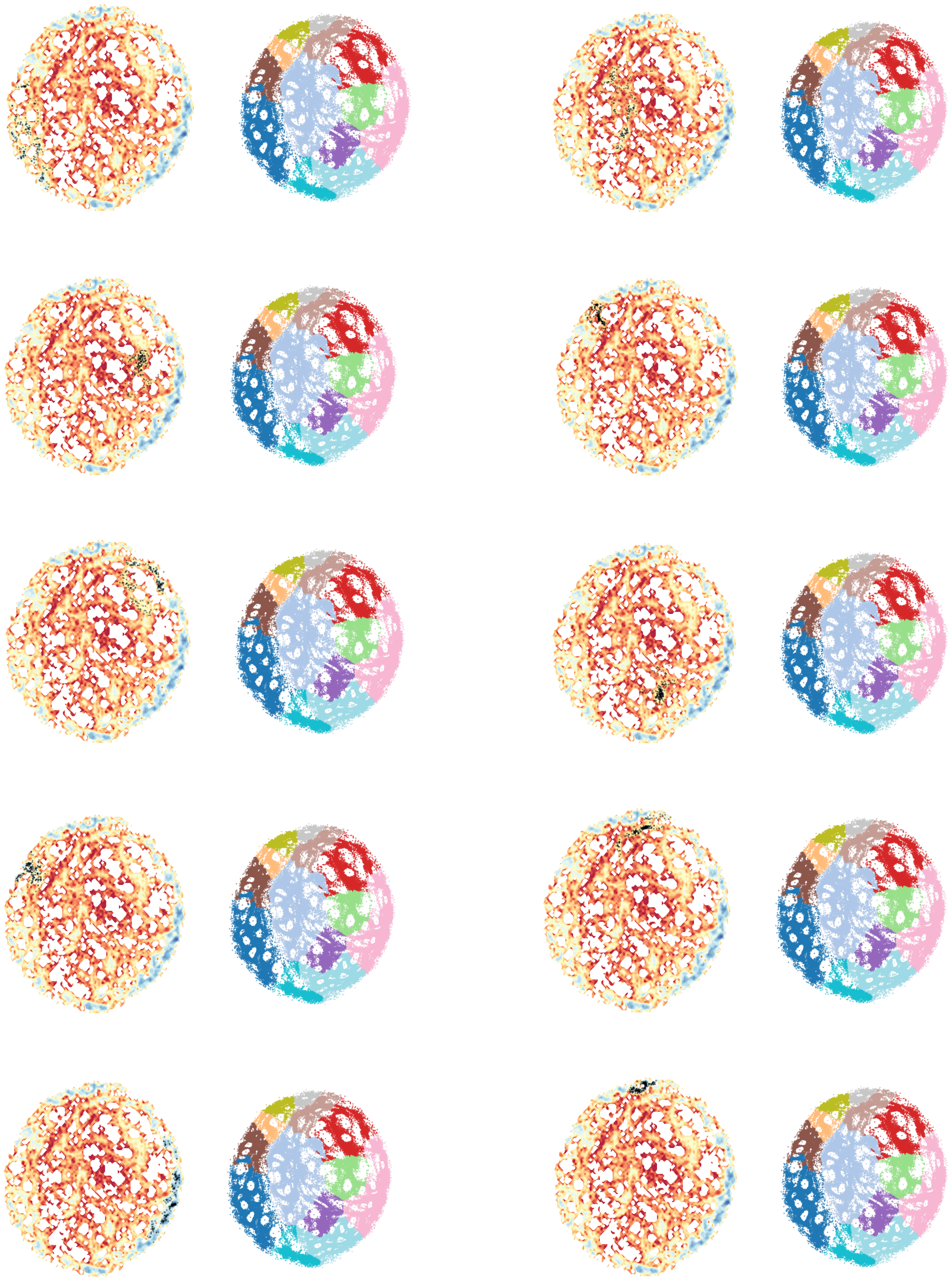

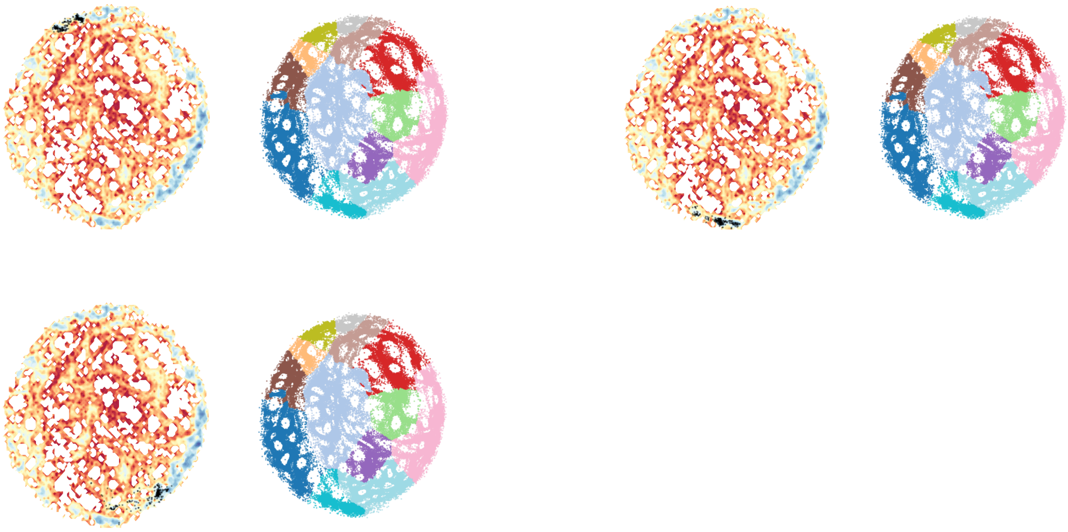

***Figure S22:*** *Single conformation extraction (black dots) for each microstate of the ELViM projection used for visualization and for structural properties (RMSD and $R_g$) characterization.*

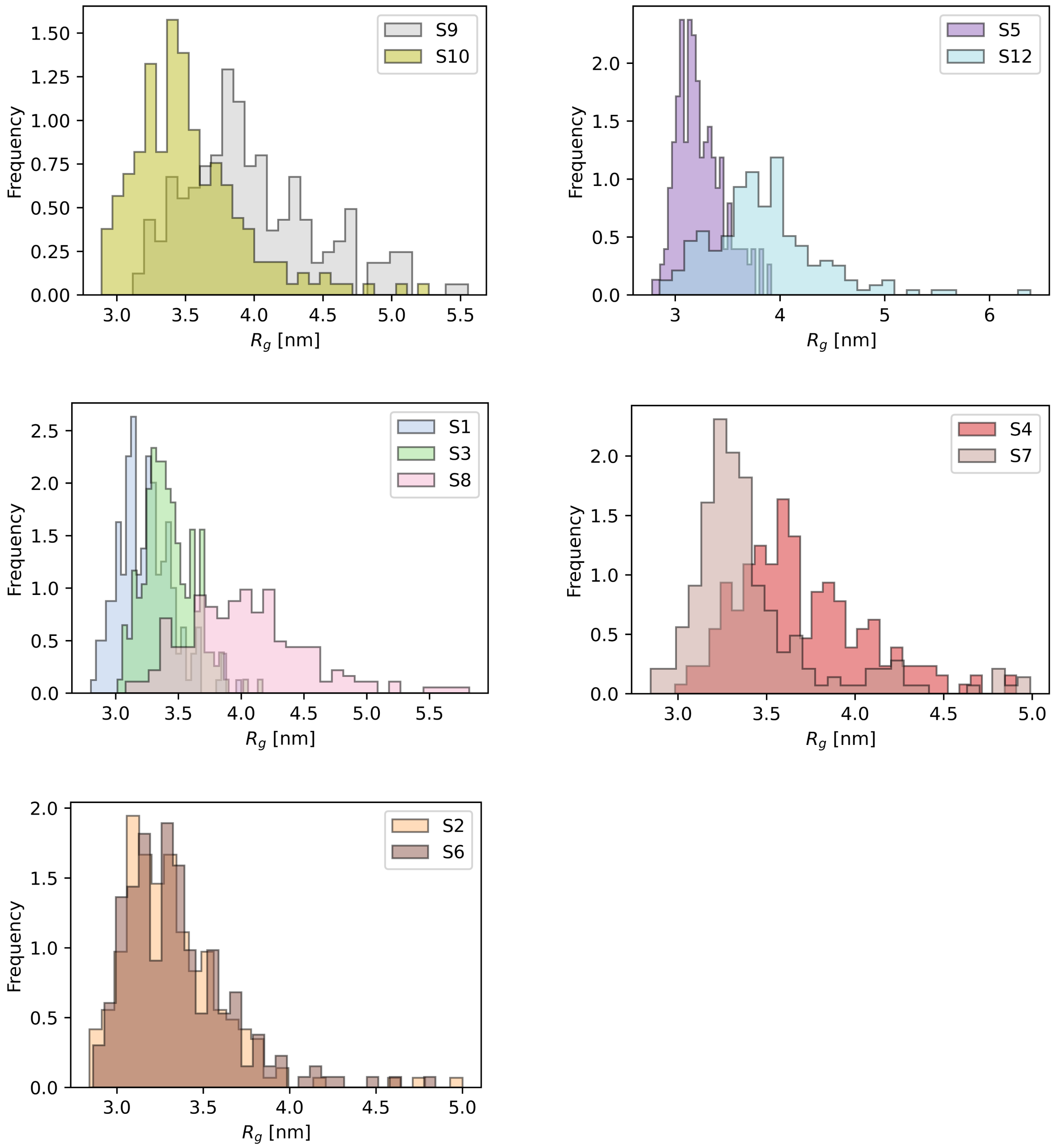


***Figure S23:*** *$R_g$ distribution comparison between states with similar domain rearrangements. Microstates showing marked differences in the radius of gyration are left separated (a,b,c,d). The last plot show very similar $R_g$ distribution but different populations in the domain rearrangements.*

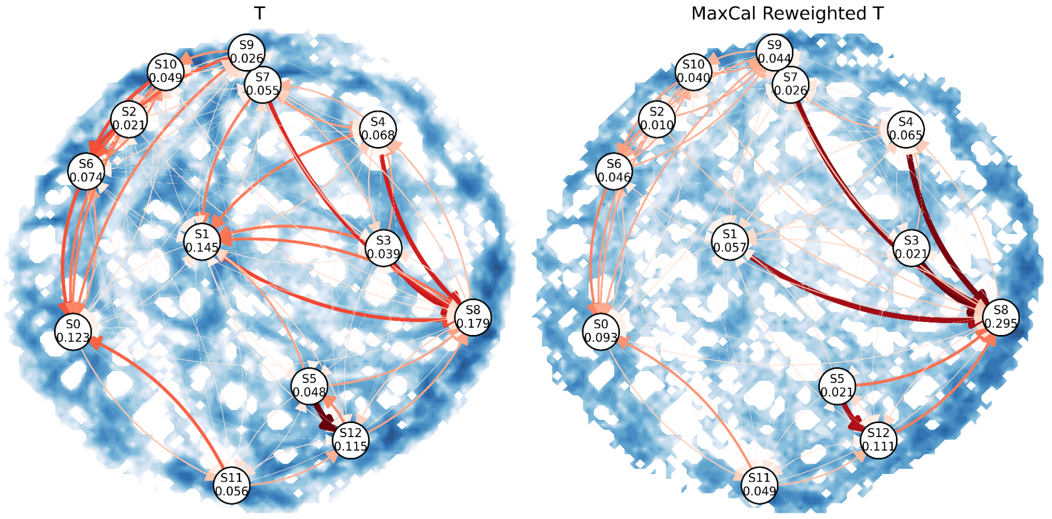


***Figure S24:*** *Transition and equilibrium probabilities obtained from the MSM on the left and the MaxCal reweighted transitions with the reweighted equilibrium probabilities on the right*

#### ***Fit of SAXS DATA***

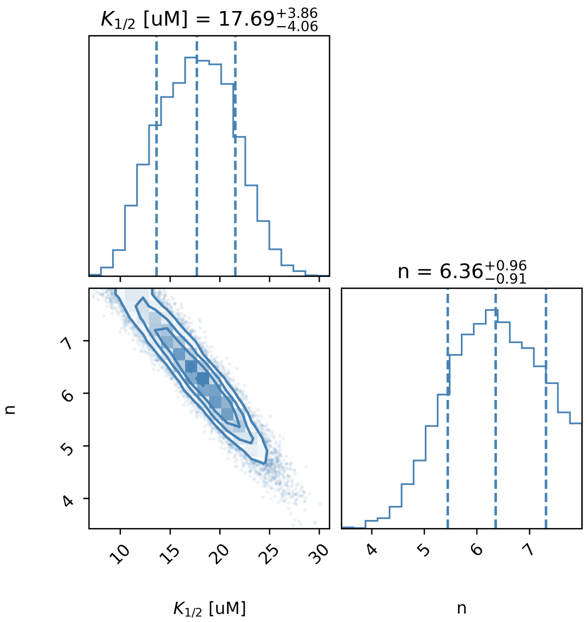


***Figure S25:*** *Corner plot of estimated parameter posterior distribution with 16th, 50th and 84th percentiles. The fit of the $R_g$ (SAXS derived) experimental values at fixed $Ca^{+2}$ concentration (50μM) and varying protein concentration [$GSN_{tot}$], yielded a half transition free calcium concentration $K_{1/2}$ = 18 ± 4μM and an effective ion-binding occupancy $n_{eff}$ = 6.4 ± 0.9. The posterior distribution sampling was done using emcee algorithm [2] assuming a likelihood with Gaussian-distributed experimental errors, with the standard deviations given by the experimental uncertainties and uniform prior distributions.*

***Table S2:*** *Predicted* $[Ca^{2+}]_{eq}$ *at different total protein concentration* $[GSN]_{tot}$

| $[GSN]_{tot}$ [μM] | $[Ca^{2+}]_{eq}$ [μM] (predicted) |
|---|---|
| 4 | 30.1 |
| 7 | 40.6 |
| 12 | 61.7 |
| 25 | 99.1 |
| 50 | 172.2 |
| 80 | 272.0 |

#### Supplementary Materials and Methods

##### 1. Protein production and SAXS analysis

For SAXS studies, full-length human GSN was produced following the protocol reported in [3]. Briefly, the protein was heterologously expressed in *E. coli* SHuffle cells (New England Biolabs) following induction with 0.5 mM IPTG and 16 h incubation at 18 °C. Cells were disrupted using a benchtop homogenizer and the clarified lysate was applied to a HisTrap HP affinity column (GE Healthcare). The sample was further purified by anion-exchange chromatography (Resource Q), followed by size-exclusion chromatography (HiLoad 16/600 Superdex 200).

The WT GSN stock was dialyzed against 20 mM HEPES, pH 7.4, 100 mM NaCl, supplemented with either 1 mM EDTA or 0.05, 0.2, 1 and 10 mM $CaCl_2$. Preliminary data that allowed to set up experimental conditions and control experiments were collected at BM29 [4] (ESRF, Grenoble France) within the MX-2691 and MX-2591 proposals. The data presented in this study were collected at B21 BioSAXS beamline of the Diamond Synchrotron (Didcot, Oxfordshire, UK) [5]. GSN at different $Ca^{2+}$ concentrations were diluted to roughly 4 mg/ml in the proper dialysis solution and further serially (1:1) diluted four more times for a total of 25 conditions (Table S1 and Figure S1).

The data analysis was performed using the program PRIMUS 3.0 (ATSAS software suite, [1]) and ScÅtter 3 (http://www.bioisis.net)).

##### 2. Design and production of double-cys variants of the protein

DEER-EPR studies required engineered GSN variants carrying pairs of solvent-exposed cysteines within the detectable distance range (< 8 nm). At the time we initiated this work, only the structure of full-length gelsolin in the closed conformation was available (3FFN, [6]), and this model was used to map all cysteine residues in the protein. Among the five cysteines present in GSN, two form a structural disulfide bond buried in the core of G2 [7], while the remaining three residues appear to be partially solvent-exposed.

To experimentally validate these predictions, we quantified free thiols using the DTNB assay under three conditions: 1 mM EDTA (closed conformation), 1 mM free $CaCl_2$ (open conformation), and 2% SDS (fully denatured). The resulting thiol/GSN ratios were 0.1, 0.5 and 3.1, respectively. These data confirm the structural hypothesis and indicate that the three free cysteines remain only partially accessible even in the active $Ca^{2+}$-bound form and thus may not be efficiently labelled.

To overcome this limitation, we removed the three partially exposed cysteines by site-directed mutagenesis (Q5® Site-Directed Mutagenesis, NEB) and reintroduced pairs of cysteines at positions predicted to be well solvent-exposed by substituting selected serines. The resulting engineered pairs were:

*N-term-G2* (C24-C226), N-*term-G3* (C24-C301), *G1-G3* (C93-C301), *G2-C-term* (C226-C742), *G3-G5* (C301-C609), *G4-G5* (C428-C609), *G4-C-term* (C428-C742).

These seven variants ensure the measurement of at least two distances per domain across all six GSN domains.

Double-cysteine variants were expressed and purified following the same procedure used for the WT protein, except that buffers used for the ion-exchange and size-exclusion chromatography steps, respectively the second and final purification stages, were supplemented with 5 mM DTT.

3. **4p-DEER (Q-band) Spectroscopy**

Spin labelling reactions on all previously listed cysteine variants were carried out using S-(1-oxyl-2,2,5,5-tetramethyl-2,5-dihydro-1H-pyrrol-3-yl)methyl methanesulfonothioate (MTSL) tag. Initially, each protein solution was incubated with a 10-fold molar excess of DTT to reduce the thiol functional group of the target residues. EDTA with 2 mM as final concentration has been added also to the solution in order to remove the presence of $Ca^{2+}$ ions from buffer solution. The mixture was kept under gentle stirring at 4°C for 30 minutes. Successively, DTT and EDTA were removed from each sample with PD-10 desalting columns using 20 mM of Hepes at pH 7.4 supplemented with 100 mM NaCl (spin labelling buffer) as eluting phase. The fractions containing GSN were pooled together, and a 10-fold molar excess of MTSL spin label was added to the solutions. The spin labelling reactions were placed at 4°C under gentle stirring. After 2 hours, a second addition of the spin label has been carried out maintaining the same reaction conditions previously described. At the end, the unreacted fraction of MTSL was removed from each protein sample using desalting PD-10 chromatography columns in the spin labelling buffer prepared in deuterated water. Cw-EPR spectra (X-band, room temperature) have been recorded to identify the fractions containing the spin labelled variants of GSN and to avoid possible contamination arising from free MTSL in solution. Then, spin labelled mutants have been concerted using AMICON centrifugal filters (MWCO = 30kDa) up to 100-120 µM, measuring the protein concentrations from the absorbance at 280 nm of the respective UV/vis spectra GSN molar extinction coefficient ε = 115530 $M^{-1}cm^{-1}$). In addition, the spin concentration of each sample has been calculated from the double integral of the cw-EPR spectrum and the relative intensity have been compared to a calibration curve. These measurements allowed us to estimate a labelling yields in the 80-90% range for all spin labelled mutants used in this study. At the end, the GSN variants have been stored at -80°C.

All the samples for 4p-DEER experiments in the absence and in the presence of different amount of calcium ion have been prepared in deuterated spin labelling buffer supplemented with 30% v/v of d8-glycerol with a final protein concentration of 80 µM. For the samples without $Ca^{2+}$, EDTA at 1 mM was also added to the solutions; for the other samples calcium ion was added to the GSN solution respectively at 0.2 µM, 2 µM, 100 µM, 200 µM and 1 mM concentrations. Samples were individually transferred into capillaries, flash frozen in liquid nitrogen, and successively placed in the cavity of EPR spectrometer.

DEER experiments have been performed on a Bruker ELEXSYS E580 operating at Q-band (34 GHz) using the EN 5107D2 resonator. The system was equipped with an Oxford helium cryostat temperature regulation unit and the data were acquired at 60 K. Four-pulse DEER experiments [8] were run with pulse durations of 20 ns (π/2) and 40 ns (π), interpulse delays $t_1$ of 200 ns and $t_2$ adjusted according to $T_m$ (typically between 2.0 to 9.0 μs). The pump frequency was centered at the maximum resonance while the observer frequency was 56 MHz lower. A 8-step phase cycling was applied to remove unwanted echoes. The total acquisition times have been between 8-16 hours to maximize signal-to-noise ratio. Background correction of the DEER traces and the distance distributions were extracted from raw experimental data with *DeerAnalysis2019* software using Tikhonov regularization [9]. Computed spin distances have been calculated using MMM2019 [10] software using PDB: 3FFN and PDB: 8VIZ respectively for the closed (absence of $Ca^{2+}$) and open (1 mM $Ca^{2+}$) states of GSN.

#### 4. SAXS fit

To extract an effective half-transition free calcium concentration, the SAXS data were fitted using an effective two-state thermodynamic model. In this model, GSN is assumed to exist in either a compact, calcium-free state (*g*) or an open state, (*g∗*), associated with the binding of an effective number of calcium ions:

$$g + n_{eff}c \rightleftharpoons g^*$$

where *c* denotes the free calcium concentration and $n_{eff}$ is the effective number of calcium ions associated with the transition. The model therefore describes the sample population as a mixture of a compact, calcium-free Gelsolin state and an open, calcium-bound state.

The effective half-transition free calcium concentration, $K_{1/2}$ is defined through the equilibrium relationship

$$K_{1/2}^{n_{eff}} = \frac{g}{g^*} c^{n_{eff}}$$

Equivalently, the population of the open state can be written as

$$g^* = g_{tot} \frac{c^{n_{eff}}}{K_{1/2}^{n_{eff}} + c^{n_{eff}}}$$

This simple two-state model was chosen to extract global parameters describing the calcium-dependent transition, rather than resolving the underlying microscopic binding mechanism. A more detailed model incorporating multiple calcium-binding and conformational states would require additional parameters and, given the number of experimental data points available, would be more susceptible to overfitting.

Within this framework, the measured radius of gyration is described as the population-weighted average of the radii of gyration of the two states,

$$R_g = R_{g,apo}\frac{g}{g_{tot}} + R_{g,1mMCa^{+2}}\frac{g^*}{g_{tot}}$$

The radii of gyration of the open and closed states, and , were determined directly from the SAXS measurements by averaging the values obtained at different GSN concentrations under calcium-saturating and calcium-free conditions, respectively.

To account for calcium depletion due to binding to GSN, the equilibrium populations were determined together with the free calcium concentration by imposing conservation of both GSN and calcium. The corresponding conservation equations are

$$g + g^* = g_{tot}$$

and

$$c + n_{eff}g^* = c_{tot}$$

where $c_{tot}$ is the total calcium concentration. Substituting the equilibrium expression for g* into the calcium conservation equation gives the following closed equation for the free calcium concentration:

$$c + n_{eff}g_{tot}\frac{c^{n_{eff}}}{K_{1/2}^{n_{eff}} + c^{n_{eff}}} = c_{tot}$$

For a given total GSN and calcium concentration, this equation was solved numerically to obtain the corresponding equilibrium free calcium concentration. The populations of the two states were then calculated as

$$g^* = g_{tot}\frac{c^{neff}}{K_{1/2}^{neff} + c^{n_{eff}}}$$

and

$$g = g_{tot} - g^*$$

The parameters $n_{eff}$ and $K_{1/2}$ were estimated using the emcee algorithm [2], which samples the posterior distribution of the model parameters given specified prior distributions and likelihood function. The likelihood was constructed assuming Gaussian-distributed experimental errors, with the standard deviations given by the experimental uncertainties. Uniform prior distributions were used for the two model parameters,

$$n_{eff} \in (1, 8]$$

and

$$K_{1/2}[\mu M] \in (0, 1000)$$

The posterior distributions of the fitted parameters, together with their 16th, 50th, and 84th percentiles, are shown in Figure S25. The resulting parameter estimates are

$$n_{eff} = 6.4 \pm 0.9$$

and

$$K_{1/2} = 18 \pm 4 \mu M$$

Using the fitted parameters, the equilibrium free calcium concentration was subsequently calculated for each GSN concentration by solving the calcium conservation equation. The predicted total calcium concentration required to reach the half-transition/free $K_{1/2}$ at each GSN concentration are reported in Table S2. The fitted parameters and the corresponding equilibrium calcium concentrations were then used to calculate the expected radius of gyration for each GSN concentration. The resulting model predictions show good agreement with the experimental data across the full range of GSN concentrations (Figure S1).

#### 5. Conventional Molecular Dynamics Simulations

Conventional atomistic, explicit solvent, MD simulations were performed to generate training data for the multi-eGO model, where interatomic interactions are obtained by combining information from conformational ensembles and prior models.

The full-length apo training ensemble was taken from our previous study [11], where the closed GSN model derived from PDB 3FFN was refined against SAXS data using metainference multireplica simulations. Ten replicas of 100 ns each were generated, and the ensemble used here corresponds to the last 50 ns of each replica, concatenated into a single 500 ns trajectory. These simulations employed DES-Amber [12] and TIP4P-D [13] in 100 mM NaCl.

MD simulations of calcium-bound single domain and calcium and actin-bound multi-domain systems (G2-G3 and G4-G5-G6) were run using the same force field and water model. Starting structures were obtained from the following PDB entries:

- G1: PDB 1RGI [14] (residues 26-149)
- G2: PDB 6QW3 [7] (residues 158-266)
- G3: PDB 6LJF [15] (residues 270-370)
- G4: PDB 1H1V/1P8X [16,17] (residues 412-535)
- G5: PDB 1H1V/1P8X [16,17] (residues 516-626)
- G6: PDB 1H1V/1P8X [16,17] (residues 635-741)
- G2-G3 actin bound: PDB 3FFK [6] (actin removed; residues 157-372)
- G4-G5-G6 actin-bound: PDB 1H1V [16] (actin removed; residues 412-741)

Domains G1 and G4 contain both type I and type II calcium-binding sites, while the remaining domains contain only type II sites.

All systems were prepared using Maestro (Schrödinger Release 2026-3: Maestro, Schrödinger, LLC, New York, NY, 2025) with protonation states assigned for pH 7.0. Each system was solvated in a cubic box using the TIP4P-D water model and neutralized with $Na^+$ and $Cl^-$ ions at 100 mM ionic strength.

All MD simulations were performed following the same protocol. The system was first energy-minimized using the steepest descent algorithm until the maximum force was below 1000 kJ $mol^{-1}$ $nm^{-1}$, followed by a conjugate gradient and an L-BFGS minimization. This was followed by a 2 ns relaxation phase with positional restraints applied to all heavy atoms, and a subsequent 500 ps unrestrained equilibration in the NPT ensemble at 300 K and 1 atm using the v-rescale thermostat [18] and the c-rescale barostat [19]. Production runs were then performed for 1 μs in the NPT ensemble at the same temperature and pressure conditions. Long-range electrostatics were calculated using Particle-mesh Ewald summation [20] with 1.0 nm Van der Waals and Coulomb cutoffs and 0.12 FFT grid spacing. A 2 fs integration timestep was used, with bond constraints applied using LINCS [21].

MD simulations were subsequently analyzed using cmdata and make_mat.py tools in the multi-eGO suite to obtain statistical information on interatomic distance distributions and contact probabilities (mode: intra and cross when calcium ions were present).

#### 6. Multi-eGO model and preparation

The multi-eGO model combines a generic description of protein conformational behavior with structure-specific interactions derived from reference ensembles. The bonded interactions define the underlying molecular topology, while the prior non-bonded interactions account for residual, sequence-independent forces that contribute to chain compaction in the absence of a stable folded structure.

Bonded terms are reoptimized for each amino-acid targeting experimental $J_{HN}^3$ coupling [22], Coil distribution extracted from high resolution PDB structures. Non-boned prior terms are parametrized to reproduce experimentally measured radii of gyration of intrinsically disordered proteins (IDPs) following the work of CALVADOS [23] or Mpipi [24]. This approach provides a transferable description of nonspecific interactions across different protein sequences and lengths.

On top of this prior model, multi-eGO introduces specific non-bonded interactions through a reweighting procedure. In this framework, a training ensemble defines the target contact statistics, while a prior ensemble provides the reference conformational distribution. The interaction strengths of the prior model are subsequently adjusted according to the difference between the contact probabilities observed in the training ($p_{ij}^{train)}$ and prior ($p_{ij}^{prior}$) ensembles, thereby enhancing or weakening specific contacts to reproduce the target structural ensemble while retaining the underlying conformational behavior of the prior model. The reweighting is defined by the following equation:

$$\varepsilon_{ij} = \varepsilon_{ij}^{prior} - \frac{\varepsilon_0 - \varepsilon_{ij}^{prior}}{\ln p_{th}} \ln\left(\frac{p_{ij}^{train}}{max\{p_{th}, p_{ij}^{prior}\}}\right)$$

A threshold probability $p_{th}$ is introduced to regularize this reweighting and prevent poorly sampled contacts from generating unrealistically strong interactions. The free parameter of the model (1 for each pair of prior-training ensemble statistics) $\varepsilon_0$ represents the maximum interaction strength.

A first Multi-eGO Prior MD simulation was performed using the molten-globule multi-eGO force-field on the full-length protein. It was performed in vacuum using stochastic Langevin dynamics 10.1021/ct3000876 using a friction time constant τ = 25 ps in the NVT ensemble for 1 µs. The simulation was then analyzed using cmdata and makemat.py in the multi-eGO suite to generate the prior matrices used to reweight intra-domain interactions (Figure S4). Interdomain multi-eGO prior simulations were performed by learning intra-domain interactions from the closed ensemble (Figure S4) using $\varepsilon_0^{intra\text{-}dom}$ = 0.28 kJ/mol to ensure folded domains. The resulting multi-eGO production force-field was used to run an additional 1 µs simulation used as the inter-domain prior model, under the same conditions used above, where domains remain folded but interact with one another only via the prior multi-eGO molten-globule interactions. The ensemble is then analyzed using cmdata and makemat.py to obtain a probability matrix which describes the contact propensity of each folded domain to interact with each other only via chain connectivity and prior interactions. These probabilities are used to reweight the interactions learned from the training ensembles in the inter-domain regions.

The final production multi-eGO force-field was constructed by merging all the information obtained from the training ensembles with the 2 priors and different ε parameters. Intra-domain interactions were learned from both the full-length apo simulation and the calcium-bound single domain simulations with $\varepsilon_0^{intra\text{-}dom}$ = 0.28 kJ/mol. Inter-molecular interactions with calcium were learned from both the holo single domain and the holo multi-domain simulations with $\varepsilon_0^{inter\text{-}mol}$= 5 kJ/mol to ensure stable binding of calcium ions modelling the effect of a saturation concentration of ions. Inter-domain interactions were learned from the full-length apo simulation and the holo multi-domain simulations. The learning of inter-domain interactions was separated into interfaces between the multi-domain subsystems (Figure S5) and the rest of the interfaces (Figure S5) learned only from the full-length closed apo training ensemble. We set the interaction energy parameter between interfaces of the multi-domain subsystems (G2-G3 and G4-G5-G6) to $\varepsilon_0^{inter\text{-}dom\text{-}1}$ = 0.16 kJ/mol. We then set the interaction between the same domain interfaces but learned from the closed apo training ensemble to $\varepsilon_0^{inter\text{-}dom\text{-}2}$ and the remaining interactions associated with all other interfaces to $\varepsilon_0^{inter\text{-}dom\text{-}far}$. We tested the different values for these last to energy scale parameters in the vicinity of 0.14 kJ/mol:

- set a): $\varepsilon_0^{inter\text{-}dom\text{-}2}$= 0.13kJ/mol, $\varepsilon_0^{inter\text{-}dom\text{-}far}$ = 0.14kJ/mol
- set b): $\varepsilon_0^{inter\text{-}dom\text{-}2}$ = 0.14kJ/mol, $\varepsilon_0^{inter\text{-}dom\text{-}far}$ = 0.14kJ/mol
- set c): $\varepsilon_0^{inter\text{-}dom\text{-}2}$=0.14kJ/mol, $\varepsilon_0^{inter\text{-}dom\text{-}far}$ =0.15kJ/mol
- set d): $\varepsilon_0^{inter\text{-}dom\text{-}2}$ =0.15kJ/mol, $\varepsilon_0^{inter\text{-}dom\text{-}far}{}_0$ =0.15kJ/mol

The overall range of values was obtained by trying to sample all the possible domain rearrangements, testing variation around the obtained values to check to which extend ensemble properties depend on the set of chosen parameters.

#### 7. SAXS calculations

SAXS profiles were computed using PEPSI-SAXS [25], by extracting frames every 500 ps after adding hydrogen atoms using GROMACS pdb2gmx [26] and interpolating the curve to the experimental q values using scipy library [27]. The agreement between the experimental and simulated SAXS profiles quantified by the reduced chi-square, is found to be χ2 ≃ 5 (Figure S6 for all the tested set of ε parameters. Mean and standard deviation of the mean of the SAXS profiles were computed over 3 replicates for each set of ε parameters.

#### 8. Bayesian Maximum Entropy Reweighting

Bayesian Maximum Entropy is a reweighting technique which finds the most probable probability distribution that is compatible with experimental data [28]. This is done by minimizing the cost function

$$\mathcal{L}(w_1 \ldots w_n) = \frac{m}{2}\chi^2(w_1 \ldots w_n) - \theta S_{\mathrm{REL}}(w_1 \ldots w_n),$$

where m is the number of experimental restraints, the reduced χ2 quantifies the agreement with experimental data and relative entropy SREL measures the deviation from the initial weights. The scaling parameter θ defines the relative confidence of the initial set of weights. The smallest the θ, the greater is going to be agreement with the experimental data at the cost of losing information from the initial data, quantified by the effective number of frames:

$$N_{eff} = exp(S_{\mathrm{REL}}).$$

We computed the BME reweighting for multiple values of the regularization parameter θ, until a χ2 ≃ 1 was reached. This value represents a good statistical agreement with experimental data without overfitting it and without losing more information from the starting ensemble than what’s necessary. For all 3 replicas and for all tested values of ε parameters, this was achieved approximately at θ = 100 as shown in figures S7. For this value of the regularization parameter, we obtain an effective number of frames Neff ≃ 70%, meaning that most of the information is retained without the need of selecting a small subset of the data.

#### 9. DEER calculations and filtering

In silico DEER profiles were computed using the DEERPREdict software [29]. The software models the SDSL paramagnetic probe explicitly using a rotamer library and computing per frame distance distribution between paramagnetic probes. After averaging over all frames, a low-pass filter is applied to the distance distribution resulting in a smoothed distance distribution P (r). The dipolar modulation signal can be backcalculated from the distance distribution, via:

$$S(t) = \int_0^\infty P(r)K(r,t)$$

where K(r, t) is the DEER kernel [29].
We calculated DEER distance distributions and dipolar signals using a low pass filter standard deviation of 0.0707 nm and maximum distance of 12 nm for all the experimental residue pairs. Initial weights used for the DEERPREdict calculation were either uniform (when no reweighting is applied) or the BME resulting weights.
The final dipolar signal and distance distribution curves and errors are then obtained as the mean and standard deviation of the mean over the 3 replicates, respectively. Comparison of the computed dipolar signal with the experimental one is obtained by computing the form factor

$$V(t) = 1 + \lambda[S(t) - 1],$$

where λ is the experimental modulation depth [30] quantifying the efficiency of the DEER pump pulse [31]. Most of the DEER signals are in good agreement with the experimental ones (Figures S11). Residue pair G3-G5 shows a marked disagreement with the *in silico* signal populating short distances compared to the experimental ones (Figure S10). To obtain a faithful ensemble compatible with experimental data, we employed a filtering scheme to assign per frame prior weights based on the agreement with G3-G5 distance profile.
Each frame prior weight is obtained as

$$w_i = exp(\eta o_i c_{DEER}(\bar{r}_i) + (1 - c_{DEER}(\bar{r}_i)))$$

where

$$o_i = \int_{r_{min}}^{r_{max}} p_i(r) p_{exp}(r) dr$$

is the overlap between the experimental distance profile (including the error) and the distance distribution per frame obtained from the rotamer distribution of the probes. The parameter η is a regularization parameter which can be thought to be related to the modulation depth (the smaller η the smaller is going to be the relative importance of the overlap), while $c_{DEER}$(r) the confidence of the DEER signal as a function of distance. In this way high confidence regions scale as exp($\eta o_i$), while low confidence regions become effectively uniform, returning to a no knowledge regime. Having no prior information on the value of η, we tested different ones (in a similar way to what is done with the theta scaling parameter in BME) as shown in Figure S10.
We chose the regularization value of η=10, balancing the final agreement and amount of information lost with the filtering. The resulting set of weights is then used as initial distribution for the BME reweighting with the SAXS data (Figures S12).

#### 10. SASA analysis

Solvent Accessible Surface Area was computed using shrake-rupley algorithm [32] implemented in MDTraj Python Library [33] using a probe with radius $r_{probe}$ = 0.6 nm. This choice of using a larger probe radius than that of water ($r_{probe} \simeq$ 0.14nm) was motivated by the objective of

quantifying the propensity of each domain interacting surface to be able to bind actin (and not simply to be solvent exposed).
The SASA was computed for each domain on those residues which are bound to the actin filament in the experimental structure (PDB 8VIZ [34]). Interacting residues were identified as those in which the relative SASA difference in presence and absence of the actin filament was greater than 20% (Figure S13).
Per domain SASA distribution compared to their values in the APO closed conformation (PDB:3FFN) [6] and actin-bound conformation (PDB:8VIZ) (without the actin filament) are shown in Figures S14. Most domains show a distribution of SASA values that is between that of the closed and the actin-bound conformation. The only exception is found to be for domain G2 (Figure S14:G2) which not only has the largest acting-interacting surface (greatest SASA) but populates values that are highly compatible with those of the actin-bound conformation.

#### 11. ELViM projection

In order to represent the ensemble of Gelsolin in the open state, given its high heterogeneity and large conformational space, we employed the Energy Landscape Visualization Method (ELViM) [35,36], which was successfully used to construct an energy landscape for Intrinsically Disordered Proteins (IDP) [37,38]. The method computes a dissimilarity matrix for each pair of structures in the trajectories and then projects them into a 2-dimentional space using a Force Scheme. We concatenated all the 3 replica trajectories into one, extracting the coordinates of every 3 Cα atom (to reduce the computational cost of the projection).
These coordinates are then given in input to ELViM to create the projection, which is then converted into a free energy by computing:

$$F(x,y) = -k_B T \log p(x,y)$$

where p(x, y) is the probability distribution computed from the projection. The reweighted FES profile was constructed in the same way by weighting each point in the projection during the construction of p(x, y) with the BME weights (Figure S17).
The Radius of gyration $R_g$ is mapped onto the projection (Figure S15), showing that expanded conformation lies in the external region of the projection. Consistently to what is expected, the effect of the reweighting tends to disfavor the more compact conformations (central part of the projection), while favoring the conformation lying in the external region, associated with more expanded conformations.
Low free energy minimum conformations are then extracted using density-based clustering HDBSCAN [39] implemented in the python scikit-learn library (Figure S16) using a minimum cluster size of 1000 and minimum samples of 200. The obtained clusters are then eventually grouped based on structural similarities of the domain rearrangements. Finally, the whole projection space is assigned to each cluster using all points membership vectors.
To further characterize the structures in each state we extracted 200 random conformations with Boltzmann probability

$$p(x_s, y_s) \propto e^{-F(x_s,y_s)/k_B T}$$

as shown in Figures S22. For each structure in each state, we computed the RMSD with respect to the closed and actin bound conformation both for domains 2-3 (PDB 3FFN,3FFK) and domains 4-5-6 (PDB 3FFN, 1P8X). A threshold of 0.55nm is used to obtain the population fraction of each state as shown in Figure 4C.
This gives a quantitative description of how much each state populates different rearrangements of the coupled domains (See Figure S23) used to define the overall state in the final representation. Where states show similar structural features in terms of domain rearrangements, radius of gyration was used to decide whether merging states or leave them distinct. Figures S23 states Rg show the differences in the distribution
of the radius of gyration of states with similar domain rearrangement, which were not merged due to the ensemble properties (different radius of gyration distributions).

#### 12. MSM and MaxCal reweighting

Using the 13 microstates obtained as explained in the previous section, we constructed a Markov-State-Model (MSM) using deeptime [40].

We first tested multiple lag time from 1 to 50 (frames) using deeptime function TransitionCountEstimator with sliding count mode obtaining then the implied timescales $t_s$ as

$$t_s = \frac{-\tau}{\ln|\lambda_i(\tau)|}$$

where $\tau$ is the lag time and $\lambda_i$ are the eigenvalues of the obtained transition matrix T. From figure S18 we observe 2 clear implied timescales which show sufficient flat distribution as a function of the lag time and a third one which reaches a plateau right before the unresolved region. To test the robustness of the Markov State Model we used the Chapman-Kolmogorov test using both 3 and 4 metastable states using multiples of lag time of $\tau$ = 25: 2,3,5,13,28 (Figure S19 and S21 respectively). We then coarse-grained the Transition matrix using pcca+ [41] and projected the eigenvectors on the microstates to understand the slow motion of the system (Figure S20). The resulting slow motions are well separated in the projection and are associated to the main domain rearrangements. In particular, the slowest motion describes the rearrangements of domain G4-G5-G6 from the closed to a disordered/actin-bound state. The second slowest motion is the detachment of domain 1 and 2 from the closed state rearrangement.
To be able to better interpret the ensemble and the transition between each microstate. We considered the weights obtained by the BME reweighting by using the Maximum Caliber Principle (MaxCal) [42] to infer the most likely change in transition probabilities of the MSM given the change in the equilibrium population of the states [43]. We adopted the same iterative approach as in the original work [43] with an error tolerance of $10^{-15}$. The result of the MaxCal reweighting is shown in Figure S24. The left panel shows the equilibrium and transition probabilities between microstates inferred from the unbiased MSM, and the right one shows the MaxCal reweighted transition probabilities after the equilibrium probabilities correction.

#### 13. Opening simulations

To study the opening process and the ordering of initial exposure of each domain we performed 50 opening simulations of Gelsolin. Simulations were performed using the multi-eGO model with the same parameters used for the final ensemble. Each simulation started from the closed conformation with type II calcium sites occupied and was initialized with random velocities extracted from Maxwell-Boltzmann distribution to ensure independent trajectories. The time of opening was defined as the first passage time for which the radius of gyration reaches the threshold of $R_g$ = 3.8nm.

$$t_{opening} = \min_t \{t : R_g(t) > 3.8nm\}$$

To characterize the temporal order of interdomain separation events during the opening transition, we computed and analyzed the distribution of separation times for each relevant domain pair.

For each domain pair, we first identified the residues forming the interdomain interface by evaluating the change in solvent-accessible surface area (SASA) upon removal of the partner domain. SASA was calculated using the Shrake–Rupley algorithm as implemented in the MDTraj Python package. Residues that are buried in the closed conformation but become solvent-exposed upon removal of the partner domain were defined as interface residues and selected for the subsequent analysis.

For each domain pair, we then calculated the distance between the mean positions of the Cα atoms belonging to the respective interface-residue sets. This distance provides a measure of the relative separation of the two domains based specifically on their interacting surfaces, rather than on their overall domain positions. Among the spatially adjacent domain pairs identified in the closed structure were G1–G2, G1–G3, G1–G6, G2–G6, G3–G6, G4–G5, and G4–G6.

The cumulative distribution function (CDF) of the separation times was then calculated for each domain pair. Two domains were considered separated when the distance between the corresponding interface centroids exceeded 3 nm. The qualitative ordering of the separation events was robust to the precise choice of this threshold, with no significant changes observed upon varying it by ±1 nm.

The G4–G5 pair was not included in the CDF analysis because these two domains remain associated throughout the opening transition. Instead, the conformational rearrangement of the G4–G5–G6 region is primarily associated with a change in the position of domain G6 in the actin-capping state.

The resulting separation-time distributions reveal a clear sequential organization of the opening process (Figure 2). The first major event is the separation of the G1–G2–G3 block from the G4–G5–G6 block, with G1–G6, G2–G6 and G3–G6 showing a very similar CDF function with a separation time of around 7ns.

This is followed by the separation of G1 from G3 (τ =35.6ns) and, subsequently, by the separation of G1–G2 and G4–G6 (τ =48.6 ns and 53.2ns respectively). Thus, the opening transition proceeds through a hierarchy of interdomain rearrangements, with the initial disruption of the interface between the two major structural blocks preceding the finer rearrangements within these blocks.

This picture is consistent with the Markov state model (MSM) analysis presented in Section S12. In particular, the rearrangements involving the G1–G2 and G4–G6 interfaces correspond to the slowest large-scale motions identified by the MSM, with these rearrangements playing a prominent role in the transition from the closed to the actin-capping state. The agreement between the separation-time analysis and the MSM therefore supports a mechanistic picture in which the opening process is initiated by separation of the two major domain blocks and is followed by slower, larger-scale rearrangements involving G1–G2 and G4–G6.
